# Sub-harmonic Entrainment of Cortical Gamma Oscillations to Deep Brain Stimulation in Parkinson’s Disease: Model Based Predictions and Validation in Three Human Subjects

**DOI:** 10.1101/2022.03.01.482549

**Authors:** James J. Sermon, Maria Olaru, Juan Anso, Stephanie Cernera, Simon Little, Maria Shcherbakova, Rafal Bogacz, Philip A. Starr, Timothy Denison, Benoit Duchet

## Abstract

**Objectives:** The exact mechanisms of deep brain stimulation (DBS) are still an active area of investigation, in spite of its clinical successes. This is due in part to the lack of understanding of the effects of stimulation on neuronal rhythms. Entrainment of brain oscillations has been hypothesised as a potential mechanism of neuromodulation. A better understanding of entrainment might further inform existing methods of continuous DBS, and help refine algorithms for adaptive methods. The purpose of this study is to develop and test a theoretical framework to predict entrainment of cortical rhythms to DBS across a wide range of stimulation parameters.

**Materials and Methods:** We fit a model of interacting neural populations to selected features characterising PD patients’ off-stimulation finely-tuned gamma rhythm recorded through electrocorticography. Using the fitted models, we predict basal ganglia DBS parameters that would result in 1:2 entrainment, a special case of sub-harmonic entrainment observed in patients and predicted by theory.

**Results:** We show that the neural circuit models fitted to patient data exhibit 1:2 entrainment when stimulation is provided across a range of stimulation parameters. Furthermore, we verify key features of the region of 1:2 entrainment in the stimulation frequency/amplitude space with follow-up recordings from the same patients, such as the loss of 1:2 entrainment above certain stimulation amplitudes.

**Conclusion:** Our results reveal that continuous, constant frequency DBS in patients may lead to nonlinear patterns of neuronal entrainment across stimulation parameters, and that these responses can be predicted by modelling. Should entrainment prove to be an important mechanism of therapeutic stimulation, our modelling framework may reduce the parameter space that clinicians must consider when programming devices for optimal benefit.

## Introduction

Deep Brain Stimulation (DBS) is a form of invasive neuromodulation, where electrical impulses are delivered to specific brain regions by implanted electrodes. In the context of Parkinson’s disease (PD), high-frequency DBS (130-180Hz) is primarily used to alleviate motor symptoms (bradykinesia, rigidity and tremor [1, 2]) when medications provide inadequate benefit. While a diverse range of effects of DBS have been observed in both behaviour and neuronal rhythms, the mechanisms underlying these responses are not fully understood.

Activity in the gamma band (approximately 30 to 100Hz) has become a target for neuromodulation as it is associated with various cognitive performance features [3] as well as motor control [4]. However, it is necessary to distinguish between broadband gamma and finely-tuned gamma (FTG). While broadband gamma reflects neuronal spiking activity over a broad frequency range (which can be as wide as 30-200Hz), FTG represents narrowband oscillatory activity with a peak frequency between 60-90Hz [5, 6]. Here, we will be focusing on FTG oscillations, which were first revealed through invasive recordings of the basal ganglia in PD patients on antiparkinsonian medications [7, 8]. These have been thought to represent a “prokinetic” brain rhythm, in contrast to “antikinetic” beta rhythms (13-30Hz). Recently, prominent FTG oscillations have been found during invasive recordings from motor cortical areas in PD [5, 9, 10], and may be associated with dyskinesias. Additionally, similar cortical oscillations have been observed in rat models of dyskinesia [11, 12].

The clinical effect of entraining FTG in patients with PD has not been directly investigated, but could be beneficial (in the absence of dyskinesia). Transcranial alternating current stimulation (tACS) at gamma frequency was observed to increase motor velocity in PD, while tACS at beta frequency saw it decrease [13]. It was hypothesised that entrainment (specifically 1:1 entrainment at stimulation frequency, as depicted in Fig 1B) of both gamma and beta oscillations would explain this observation by enhancing “prokinetic” and “antikinetic” rhythms, respectively. Additionally, a shifted FTG peak frequency has been noted in the motor cortex in response to high-frequency DBS of the STN [5, 14, 10]. The gamma peak, off-stimulation between 65 and 80Hz, locks to the half harmonic of stimulation (see Fig 1), corresponding to 1:2 entrainment. Together, these studies suggests that the entrainment of FTG by DBS could play a role in ameliorating PD-associated motor symptoms. Moreover, the half harmonic lock indicates that the response to stimulation is complex and goes beyond entraining rhythms solely at the frequency of stimulation or suppressing them. However, there is currently no theoretical understanding of 1:2 gamma entrainment in PD and generally no framework to predict the occurrence of specific entrainment regimes in response to brain stimulation.

**Figure 1:**
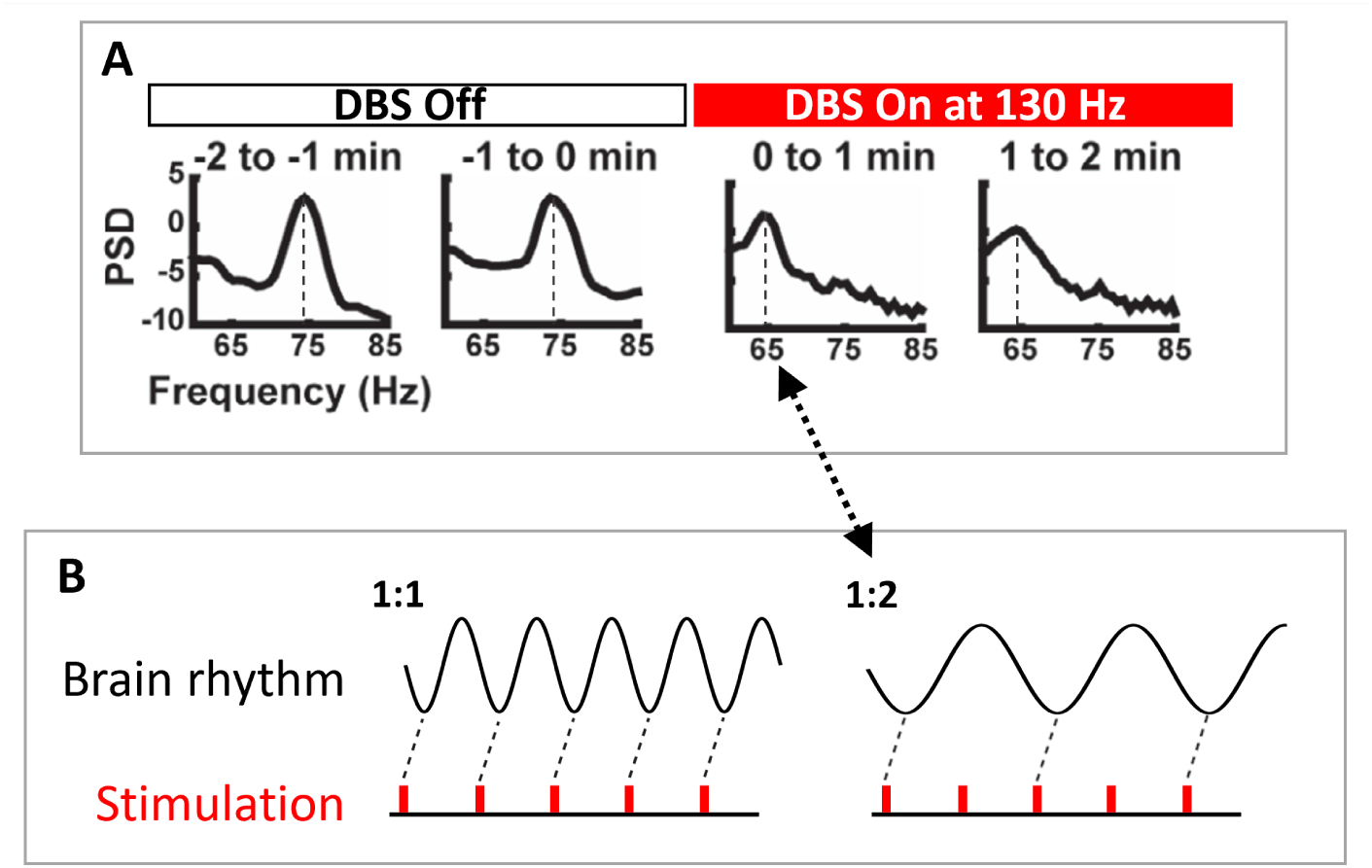
Prior human recordings demonstrate 1:2 entrainment of cortical gamma rhythms from subcortical stimulation. (A) PSD of gamma band activity recorded from the motor cortex before and during DBS to the STN at 130Hz. In the DBS Off state, a natural *∼*75Hz gamma rhythm can be observed. This is entrained at 65Hz during the following two minutes of DBS On at 130Hz. (B) 1:1 entrainment corresponds to one stimulation pulse per brain rhythm cycle and a rotation number of 1, 1:2 entrainment corresponds to two stimulation pulses per brain rhythm cycle and a rotation number of 0.5. Hence, during 1:2 entrainment, the brain rhythm locks to a frequency of half that from the external stimulation. This corresponds to the DBS On state of panel A. Panel A is adapted from [5] (with no permission required).

In this study, we use a model-based approach that utilises chronic invasive cortical recordings in PD patients to predict the properties of entrainment of cortical activity by basal ganglia DBS. We postulate that by constraining the parameters of a neuronal population model, it will be possible to predict stimulation parameters that lead to 1:2 gamma entrainment for patients with off-stimulation (medication induced) FTG. We provide a theoretical introduction to 1:2 gamma entrainment through the concepts of Arnold tongues (regions of entrainment in the stimulation frequency and amplitude space) and rotation number (the ratio of the average number of oscillation cycles to stimulation pulses). We proceed to develop a patient-specific approach by fitting a model representing interacting neural populations, the Wilson-Cowan model, to features of invasive chronic electrocorticography (ECoG) data recorded off stimulation from patients with PD. Using the fitted-models, we predict the regions of 1:2 entrainment in the stimulation parameter (frequency and amplitude) space. We proceed to verify key features of the 1:2 entrainment regions with follow-up recordings from the same patients. Lastly, these results are discussed and the implications are highlighted for future stimulation therapies.

## Materials and methods

### Arnold Tongues and Rotation Number

The frequency locking (entrainment) behaviour of a neural rhythm to external stimulation across stimulation frequency and amplitude can be described by Arnold tongues [15]. Arnold tongues are the regions in the stimulation frequency and amplitude space where frequency-locking occurs. Frequency locking is observed when a rotation number of the form p:q, where p and q are coprime integers, is maintained for several stimulation periods. In general, the rotation number may not be a ratio of integers (it is a ratio of integers only when there is frequency locking), and corresponds to the average number of oscillatory cycles achieved by the rhythm between two periodic pulses of the driving stimulation. This is calculated for periodic signals as

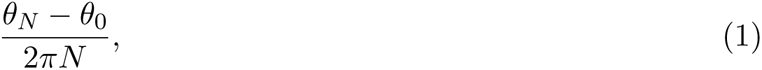

where *θ_N_* is the phase after N stimulation pulses (in this study, N *>* 50) and *θ*_0_ is the initial phase. Previously, Arnold tongues have been used to describe 1:1 entrainment in response to noninvasive neuromodulation [16, 17, 18, 19]. Depending on the system considered and the stimulation waveform, Arnold tongues can theoretically exist for various rotation numbers, including p:q with large p and/or q. However, in real systems, often only the tongues corresponding to the most stable rotation numbers, with low p and q values, will be observed. Arnold tongues often have different shapes for different dynamical systems. Generally, an Arnold tongue expands in width across larger frequency ranges as stimulation amplitude increases. This continues up until an amplitude where the tongue may border a region with another frequency-locking ratio or lose entrainment altogether.

The model we use to predict 1:2 entrainment based on off-stimulation recordings only is introduced next. How Arnold tongues are obtained in this model is described later in the *Providing Stimulation and Entrainment Analysis in the Model* section.

### Wilson-Cowan Model

The Wilson-Cowan model is well-suited to fit population-level brain recordings. The Wilson-Cowan model is a mean-field model describing interacting neuronal populations [20, 21] and, hence, is a natural choice to represent off-stimulation ECoG recordings and predict 1:2 gamma entrainment. While the sine circle map, a simpler model, can provide a first level description of 1:2 gamma entrainment (see Supplementary Materials section C), it only represents a single neural oscillator and cannot fit population-level brain recordings. The Wilson-Cowan model has been used in the analysis of neuronal responses to periodic and varying stimulation [22, 23, 24, 25, 26] and in theoretical studies of entrainment [27, 28]. Additionally, the model has been used in the analysis of resonances [29], as well as in the communication of information [30]. The Wilson-Cowan model has a limited number of model parameters which makes it feasible to constrain the model with a low risk of over-fitting the data. Despite the relatively small number of parameters, it is also able to capture a wide variety of dynamics [31, 32, 27].

We use the two-population Wilson-Cowan model to represent excitatory and inhibitory cortical populations with reciprocal connections (see Fig 2). We do not include subcortical populations in this model, and assume that subcortical stimulation is transmitted unperturbed to the cortex. The Wilson-Cowan model can be used to predict the interactions of large groups of neurons and outputs the activity of an excitatory and an inhibitory population. The population activities are denoted by *E* and *I*, respectively, and are proportional to the firing rate of that population’s neurons. Stochastic differential equations describe the evolution of *E* and *I* as

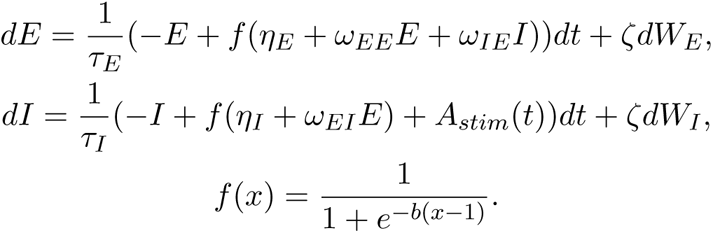

**Figure 2:**
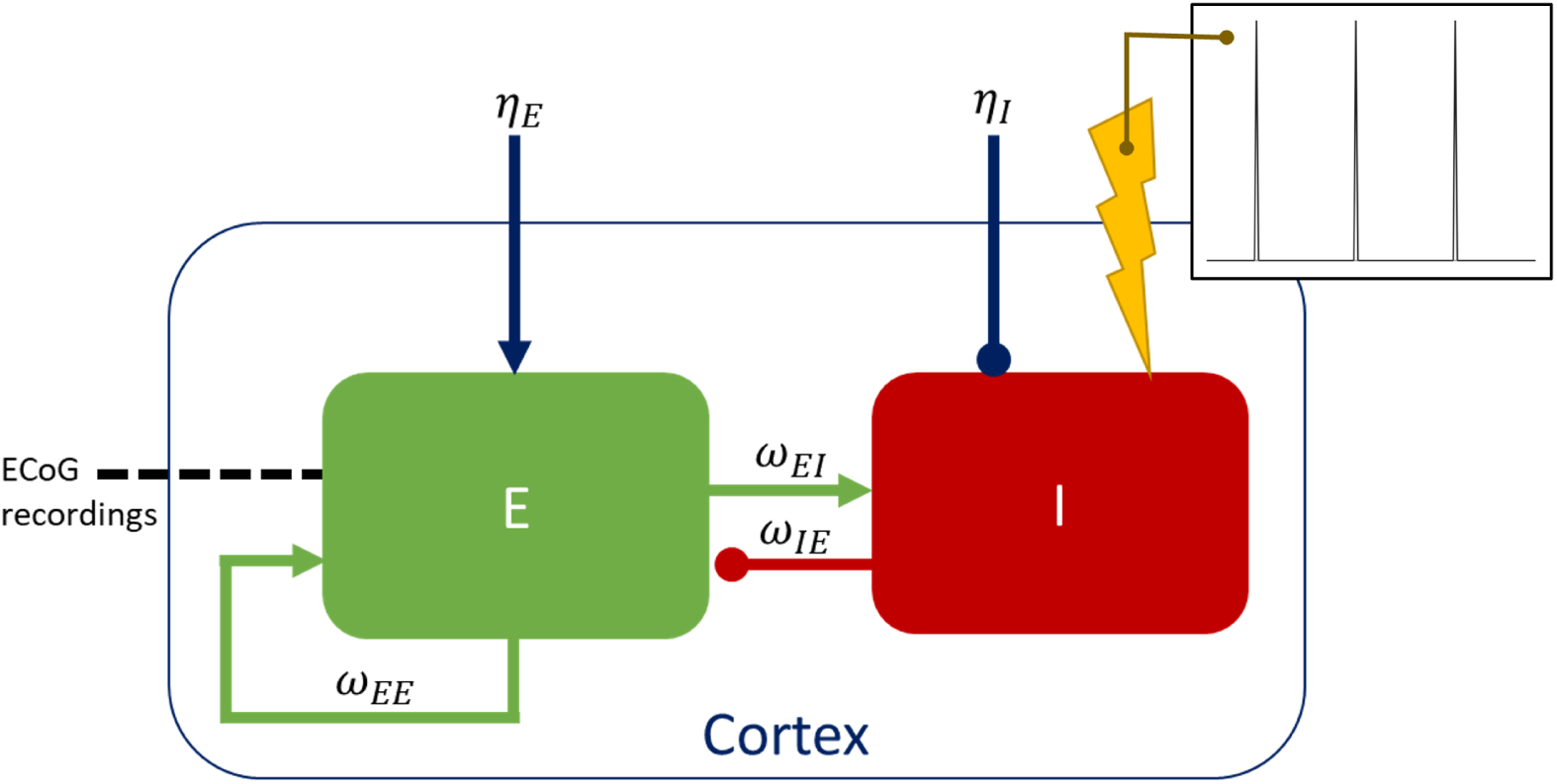
The two-population Wilson-Cowan model. Stimulation is applied to the inhibitory population (I) and data recorded from the excitatory population (E). The weights of the three connections present in this model are *ω_EI_* (excitatory connection from *E* to *I*), *ω_IE_* (inhibitory connection from *I* to *E*), and *ω_EE_* (self-excitatory connection from *E* to *E*). Additionally, there are external inputs, *η_E_* and *η_I_*, to each population. The insert displays the single time step stimulation pulse with no recharge used throughout this study (pulses with different active recharge durations, and with passive recharge are considered in Supplementary Materials section D.2).

These interactions are weighted by coupling strengths denoted *ω_P Q_* (going from population *P* to population *Q*), and occur through a sigmoid function, *f* (*x*), of steepness coefficient *b*. *τ_E_* and *τ_I_*represent the time constants of the excitatory and inhibitory populations, respectively. *η_E_* and *η_I_* are the constant inputs to the respective populations. Stochasticity is introduced to the model through Wiener processes, *W_E_*and *W_I_*, with noise standard deviation denoted by *ζ*. Noise is required to reproduce the off-stimulation data, which is characterised by bursts of activity rather than perfectly periodic dynamics (see Fig 5D).

It is unclear whether the effect of subcortical stimulation on the motor cortex is excitatory or inhibitory (more details in the Discussion section). In this study of the cortical response, we model the effect as inhibitory by applying periodic high-frequency stimulation, *A_stim_*(*t*), to the inhibitory population, while the ECoG local field potential (LFP) is modelled as the activity of the excitatory population. However, we also verify in Supplementary Materials section D.5 that our results hold for stimulation applied to the excitatory population and ECoG LFP obtained as the activity of the inhibitory population. Additionally, stimulation is applied directly to the inhibitory population, not through the sigmoid function, as this provides a greater wealth of dynamics by avoiding saturation effects [26].

### Data Collection

Human neural data were collected from three patients with Parkinson’s disease (Table 1). Cortical data off-stimulation were collected to fit the Wilson-Cowan model. On-stimulation data (subcortical stimulation) at variable stimulation frequencies and amplitudes were then used to validate predictions from the fitted model.

Patients were selected for participation in this study based on the presence of a peak in gamma band activity in the primary motor cortex off-stimulation (which is a necessary condition to fit a model to off-stimulation gamma activity). Patients were diagnosed with idiopathic Parkinson’s disease by a movement disorders neurologist and underwent DBS surgery of either the subthalamic nucleus (STN) or globus pallidus (GP). Target choice was based on the patients’ neuropsychological test results, which indicated pallidal implantation for patients with mild cognitive impairment or history of clinical depression [33, 34]. Patients were bilaterally implanted with the Medtronic Summit RC+S bidirectional neural interface (clinicaltrials.gov identifier NCT03582891, USA FDA investigational device exemption number 180097, IRB number 18-24454) quadripolar cylindrical leads into subcortical nuclei (Medtronic model 3389 or 3387 for STN and pallidum, respectively), and subdural paddle-type leads over the primary motor cortex in the subdural space (Medtronic model 0913025), see Figs 3A1-3 for RCS02, Figs 3B1-3 for RCS10 and Figs 3C1-2 for RCS18. Implantations of subcortical leads were performed using frame-based stereotaxy, and confirmed by intraoperative cone beam CT and microelectrode recording (MER) in the awake state using standard methods in RCS02 and RCS10 [35], and by intraoperative cone-beam CT alone in RCS18 [36]. For RCS10, the active contact array was localised in the GP using MER mapping of single-unit cells to traverse the postero-lateral regions of the external globus pallidus (GPe) and globus pallidus internus (GPi), Fig 3B1. For RCS02, the motor territory of the STN was confirmed by eliciting movement-related single-cell discharge patterns during MER when the DBS lead was placed with the middle two contacts in dorsal STN. Surgical placement of subdural cortical paddles has been previously described in detail [37]. Similar to subcortical contacts, localisation of subdural paddles was confirmed by computationally fusing a postoperative CT scan to the preoperative planning MRI scan, Figs 3B1-3, A2-3 and C1-2.

**Figure 3:**
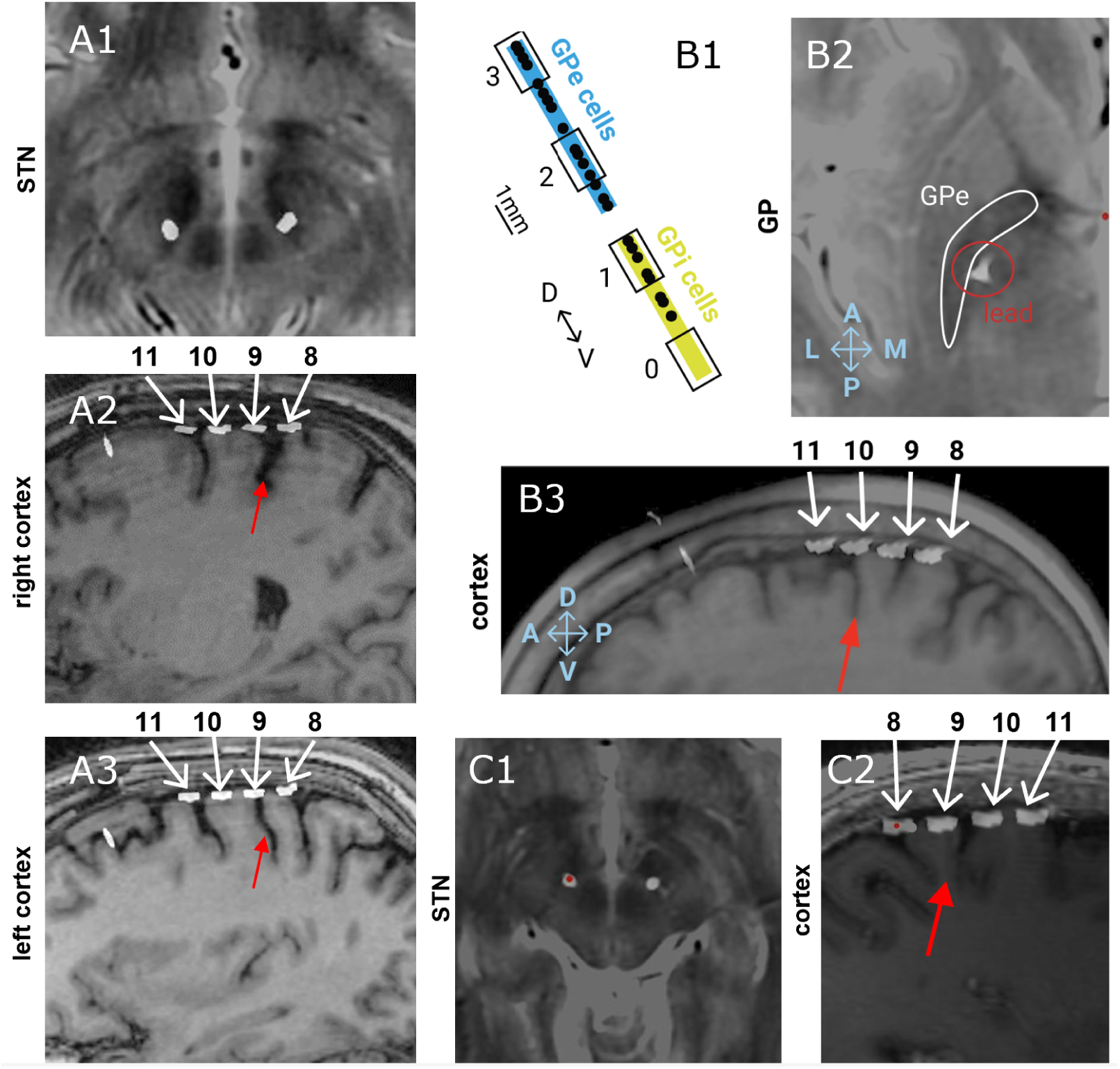
Contact localisation across the three subjects. The patients are presented as RCS02 (A1-3), RCS10 (B1-3) and RCS18 (C1 and 2). (A1-3, B2-3, C1-2) Localisation of contacts with a postoperative CT scan that is computationally fused with the preoperative planning MRI scan.(A1 and C1) subthalamic nucleus (STN) lead on a T2-weighted MRI scan. In C1, the red dot indicates the left STN lead. (A2-3, B3, C2) Quadripolar subdural paddle lead on sagittal T1-weighted MRI shows the relationship between the central sulcus (red arrow) and contacts (white numbered arrows). (B1) globus pallidus (GP) contact localisation (black numbered rectangles) with respect to the boundaries of the internal globus pallidus (GPi) (yellow) and external globus pallidus (GPe) (blue) as defined by micro-electrode recording mapping of single-unit cells (black dots). (B2) GP lead on an axial T2-weighted MRI, which visualises the GP as regions of T2 hypointensity (GPe highlighted by a white contour).

Prior to the initiation of standard therapeutic DBS, we recorded four-channel LFPs of the cortical, and pallidal (n=1) or subthalamic (n=2) sites of each hemisphere between two and four weeks post implantation. The data were streamed from patients during normal activities of daily living and while on their schedule of antiparkinsonian medication. Only the cortical prestimulation data were used to fit the Wilson-Cowan model. The recording methods and data processing were similar to those described in Gilron et al [10]. Briefly, neural data were recorded from subcortical (STN or GP, as per patient’s DBS target; Fig 3A1, B2, C1) and cortical (contacts 11-10 and 9-8 for all patients; Fig 3A2-3, B3, C2) sensing electrodes via a patient-facing graphical user interface application. Neural data were collected at 250 Hz. The Summit RC+S has two onboard filters applied after digitisation that were set to a high pass of 0.85 Hz and a low pass of 450 Hz before amplification followed by 1700 Hz low pass filter after amplification.

After several months of continuous subthalamic (n=2) or pallidal (n=1) stimulation at clinically optimised parameters, we conducted a follow-up in-clinic recording session with each subject in their on-medication state to validate the initial model predictions. During recordings, we cycled through a range of stimulation frequencies and amplitudes to further explore the DBS parameter space (Table 1). In each participant, we delivered stimulation at their clinical contact in the STN or GP, while recording from cortical sensing contacts 11-10 and 9-8. In each trial, we tested a single frequency-amplitude combination for 30 seconds while the patient was at rest. The number of data points collected was different in each patient due to patient fatigue. Filter settings for cortical contacts were the same as during off-stimulation recordings.

**Table 1:** Patient information summary. LEDD: levodopa equivalent daily dose, UPDRS: Movement Disorders Society Unified Parkinson’s Disease Rating Scale, STN: subthalamic nucleus. “Time of charting” refers to on-stimulation data collection to test model predictions. *RCS18s right lead was deactivated during testing since it was recently reimplanted and undergoing clinical optimisation.

| Patient | RCS02 | RCS10 | RCS18 |
| --- | --- | --- | --- |
| Age (years) | 54 | 64 | 67 |
| Disease Duration (years) | 7 | 13 | 7 |
| Gender | M | F | M |
| LEDD (mg/day) | 1425 | 1083 | 2031 |
| UPDRS-III off-med | 49 | 89 | 49 |
| UPDRS (off-on) | 90% | 53% | 76% |
| DBS Target | STN | Pallidum | STN |
| Duration of chronic DBS prior to collection of entrainment data (months) | 30 | 10 | 7 |
| Clinical Stimulation Settings | R: 1-C+, 2.6 mA, 60 us, 130 Hz L: 2-C+, 2.4 mA, 60 us, 130 Hz | R: 1-2-C+, 5.0 mA, 90 us, 150 Hz L: 2-C+, 5.0 mA, 90 us, 150 Hz | R: 2-C+, 2.3 mA, 60 us, 130 Hz L: 2-C+, 2.4 mA, 60 us, 130 Hz |
| Titration Ranges | R/L: Freq.: 110-170 Hz, Amp.: 0.7-3.1 mA | R/L: Freq.: 130-150 Hz, Amp.: 0-6.5 mA | R: OFF* L: Freq.: 40-180 Hz, Amp.: 0.9-4.3 mA |

### Fitting Process

To fit the parameters of our Wilson-Cowan model to prestimulation cortical FTG, we processed the off-stimulation cortical recordings (over three hours for each subject, collected between two and four weeks post implantation) to obtain data features for each patient. We separated the off-stimulation sessions into epochs with a minimum of 30 seconds of continuous and uninterrupted recordings. For the fitting process, we only used one epoch for each patient which was selected by identifying the epoch with the most prominent cortical gamma peak within the frequency range *±*3Hz of the approximate average peak gamma frequency of the overall dataset (75Hz for RCS02 and RCS10, 78Hz for RCS18). From this epoch, the signal was band-passed between *±*3Hz of the gamma peak and three features were selected for the purposes of fitting the model to each patient; the power spectral density (PSD) of the cortical FTG signal, its envelope PSD, and its envelope probability density function (PDF). These features are shown for RCS10 in Fig 4A1-3, and for RCS02 and RCS18 in S1 Fig S.2 in Supplementary Materials. The envelope is the modulus of the analytical signal of the band-passed recording and refers to a curve that traces the upper bound of the signal, providing a measure of the oscillation’s amplitude. The envelope PDF represents the distribution of the values of the envelope over the duration of the signal. These features were selected to provide a representation of the signal and its envelope in the frequency domain, as well as a representation of the statistics of the envelope in the time domain. We demonstrate in Supplementary Materials section A.1 that there is little correlation between the three features mentioned here, and that all three features are required to capture the full dynamics of the data. Fitting to off-stimulation features ensured that any presence of 1:2 entrainment is not predetermined, as could have been the case if fitting to on-stimulation data.

**Figure 4:**
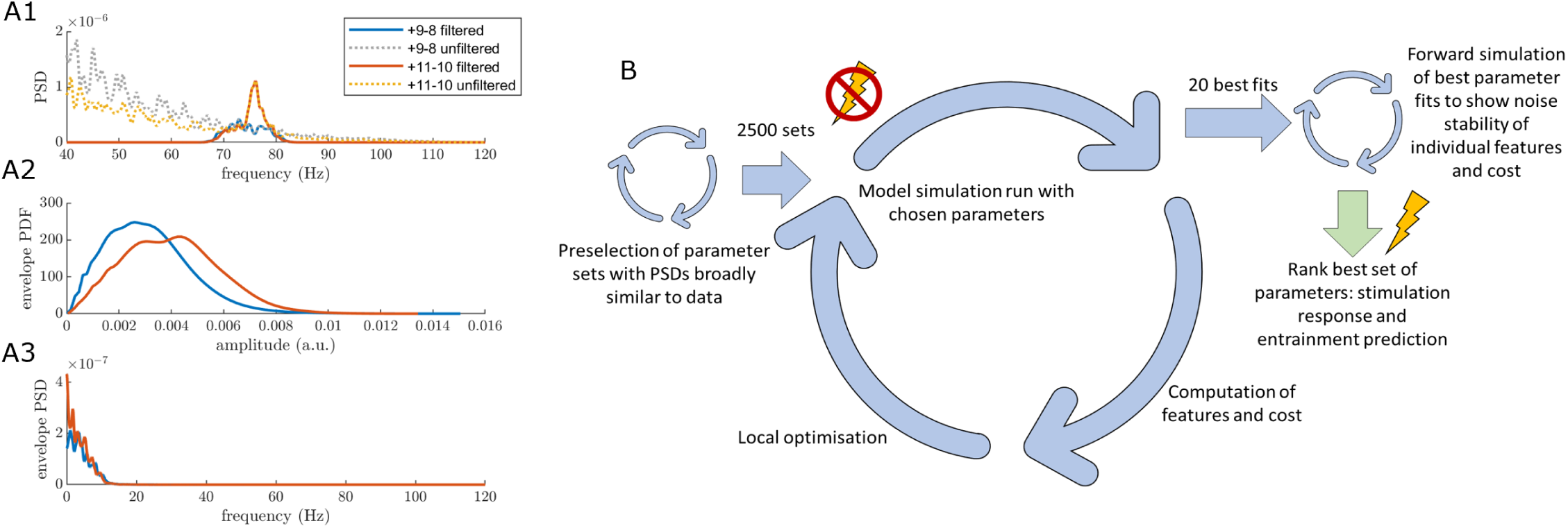
Use of prestimulation human cortical recordings to fit Wilson-Cowan model parameters. (A1-3) The three data features from RCS10 cortical recordings: power spectral density (PSD) (A1), envelope probability density function (PDF) (A2) and envelope PSD (A3), for the selected epoch, based on the gamma peak height in the cortical 9-8 and 11-10 contact. The features shown are from the cortical contacts as labelled in 3B3. The solid orange and blue lines display the band-pass filtered cortical signals between 72Hz and 78Hz, as the peak occurred at 75Hz for this patient. The yellow and grey dotted line in the PSD plot show the unfiltered signal which, for the 11-10 contact, still displays the FTG peak seen in the filtered data. The fitting is based off the filtered data from the 11-10 contact. (B) The optimisation pathway for fitting the model to off-stimulation data. This process is broken down into three main loops, as discussed in the *Fitting Process* section. Once a fitted set of model parameters is obtained we are able to make predictions for the neuronal population responses in the on-stimulation state.

**Figure 5:**
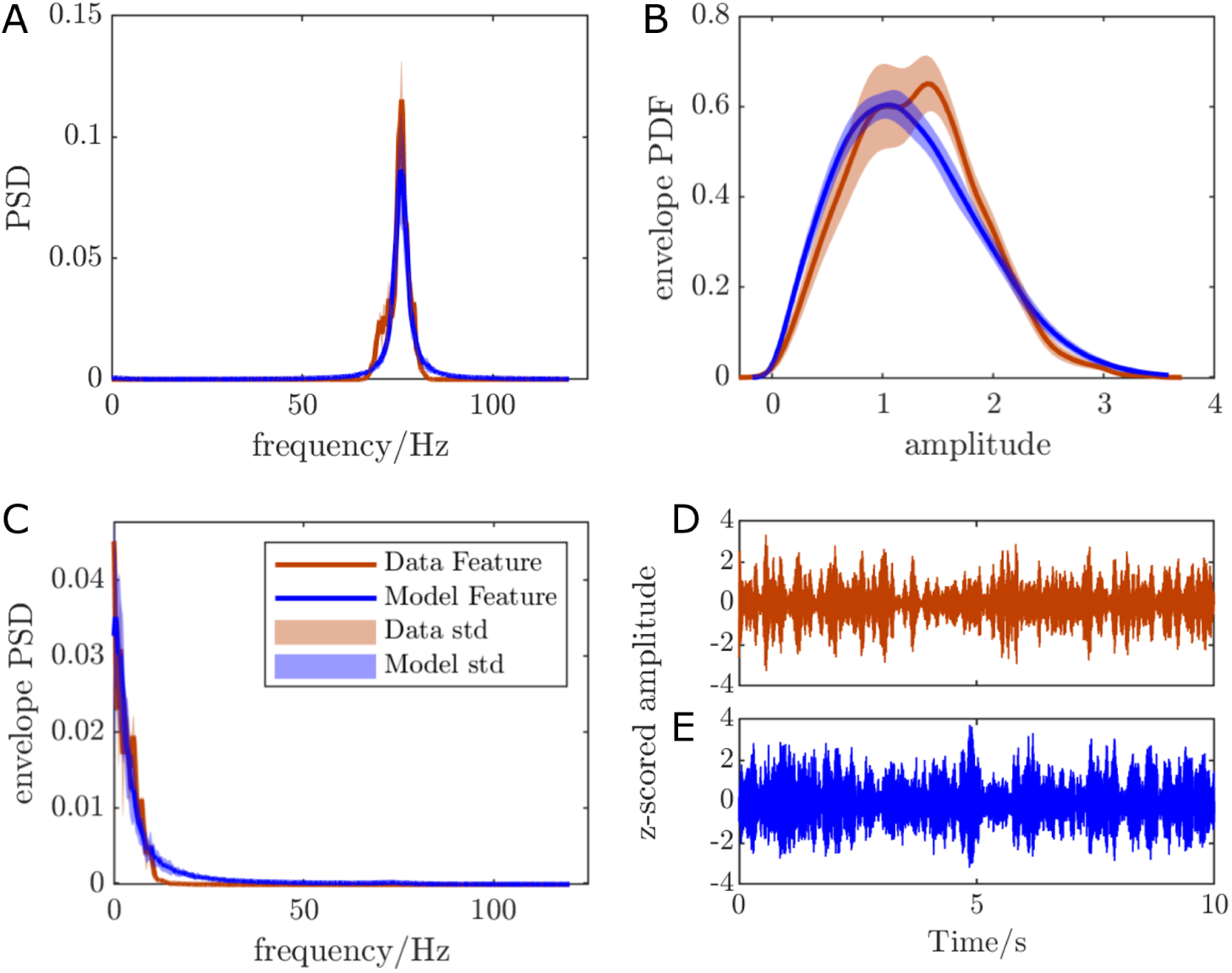
Comparison of the cortical FTG features from z-scored data and the features from z-scored model output of the best ranked model parameter set for RCS10. *R*^2^ = 0.944 on average across 50 simulations lasting 100 seconds each. The model closely matches the data PSD (A), envelope PDF (B), and envelope PSD (C). (D-E) Comparison of the band-passed off-stimulation time series from patient data (D) and model output (E) show that the model is capable of replicating similar time-domain dynamics.

Model parameters were then optimised to best match the selected cortical FTG data features (Fig 4A1-3). This process follows a fitting methodology similar to [26, 38], repeated for each patient. It begins by generating random sets of parameters, and selecting parameter sets with a PSD broadly similar to that of the data (the first loop of Fig 4B), i.e. with a gamma peak between 70 to 80Hz. This improves the computational efficiency of the parameter fitting. Accepted parameters enter an optimisation loop (the second loop of Fig 4B) using the patternsearch function of Matlab2020b, which minimises the cost function capturing the distance between model and data features (see Supplementary Materials section A.2). We run this optimisation to obtain approximately 2500 parameter sets fitted to the 30 seconds of off-stimulation cortical data, each corresponding to a different local minimum of the cost function. From the resulting fits, we select the 20 with the greatest *R*^2^ values and perform further simulations to refine the ranking of the fits (the third loop of Fig 4B), more details can be found in Supplementary Materials section A.3. The top-ranked fit is then selected based on these simulations. We don’t expect overfitting to be an issue given that we are fitting to off stimulation data, where there is no entrainment in the signal. Predictions of the response to external stimuli are then made by introducing stimulation to the off-stimulation fitted model.

### Providing Stimulation and Entrainment Analysis in the Model

We obtain entrainment predicitons in the model (i.e. Arnold tongues, in particular the 1:2 Arnold tongue) by providing high-frequency DBS at variable stimulation amplitudes and frequencies. The stimulation pulse provided throughout the majority of the modelling work in this study, unless mentioned otherwise, is a single time step positive pulse with no recharge (see the insert in Fig 2). This stimulation pulse was chosen for simplicity. Different recharge lengths and stimulation waveforms are explored in Supplementary Materials sections D.2 and D.3. Stimulation is introduced in the model as described in the *Wilson-Cowan Model* section.

In the presence of stimulation, entrainment is assessed in the model by computing the rotation number using equation (1) where *θ_i_* is taken as the unwrapped Hilbert phase of the excitatory population activity after *i* stimulation pulses. This was calculated over 50 stimulation cycles and averaged over five repeats at each stimulation parameter. To compare with data, the PSD of the model output with stimulation applied was calculated using Welch’s PSD estimate over the same number of stimulation cycles and repeats as the rotation number. The peak PSD was calculated as the maximum power in the 0 to 200Hz frequency range.

### Entrainment Analysis of the ECoG Recordings

Entrainment of cortical FTG to DBS was assessed in the data using a spectral method since the rotation number cannot be reliably computed in the data. The PSD of the data (from the cortical 9-11 contact in RCS02 and 11-10 contact in RCS10 and RCS18) was calculated in a similar way to that of the model, using Welch’s PSD estimate. Only frequencies recorded within the 40 to 120Hz range were considered when computing the peak power. To determine whether each set of stimulation parameters achieved entrainment we first determined the approximate PSD slope on stimulation by averaging the PSD in the ranges [*f_s_/*2 *−* 10*, f_s_/*2 *−* 5] and [*f_s_/*2 + 5*, f_s_/*2 + 10], where *f_s_* denotes stimulation frequency. These ranges were chosen to avoid spectral changes around the half harmonic. The signal was then determined to be showing 1:2 entrainment if the height of the PSD at half the stimulation frequency was over three times greater than the approximated baseline expected from the measured slope. This placed the threshold over any variation due to noise.

### Correspondence Between Stimulation Amplitude in the Model and in the Data

While the stimulation frequency unit is the same in the model and the data (Hz), the stimulation amplitude unit is arbitrary in the model. To correlate peak PSD predictions in the model and the data, it is therefore a pre-requisite to establish a correspondence between stimulation amplitude in the model and in the data. This was done by selecting the scaling of the model stimulation amplitude axis that maximised the correlation of peak PSD of 1:2 entrained points (as assessed in the data) between the data and the model (grid search over a linear scale).

Optimising the scaling of the model stimulation amplitude introduces an additional free parameter which needs to be accounted for in statistical analyses. For each patient, significance of the correlation between peak PSD predictions in the model and the data is therefore assessed using an approximation of the distribution of correlation coefficients under the null hypothesis (no correlation) obtained from 10,000 permutations. For each patient, we randomly reassign PSDs from the data to the different stimulation parameters used. Each permutation is then compared against model PSD predictions and the scaling is optimised to maximise the correlation between the peak PSD at the half harmonic of stimulation frequency between the model and the data, as outlined above. Only permutations which resulted in at least five 1:2 entrained results from the model simulations were accepted. P-values are then calculated using the resulting (approximately) normal distribution of correlation coefficients.

## Results

We fitted the Wilson-Cowan model, a model of interacting neuronal populations, to prestimualtion ECoG recordings in three patients with PD. Using the fitted models, we predicted for each patient the stimulation parameters leading to 1:2 entrainment of FTG. We then verified these predictions in follow-up on-stimulation recordings from the same patients.

### Prediction of 1:2 Entrainment of FTG Using a Fitted Wilson-Cowan Model

For all patients, the Wilson-Cowan model fitted to the patient’s prestimulation cortical FTG successfully reproduced all three FTG features.The top ranked model parameter sets (Table A in Supplementary Materials) had average *R*^2^ values (across 50 simulations) of 0.961, 0.944 and 0.868 for RCS02, RCS10 and RCS18 respectively, with good or satisfactory fits across all three features (Fig 5 for RCS10, and Fig S.3 in Supplementary Materials for RCS02 and RCS18). The fitted models oscillate at the patient’s FTG natural frequency (between 75-80Hz in the absence of stimulation), as seen in Fig 5A for RCS10, and Fig S.3A1 and B1 in Supplementary Materials for RCS02 and RCS18. In the presence of low amplitude stimulation, all three models display a 1:2 Arnold tongue (i.e. entrainment region) around a stimulation frequency between 150-160Hz (the 1:2 tongue stems from twice the natural frequency of the model). This is shown by the green 1:2 Arnold tongue in Figs 6A, D and G, which correspond to a region of 1:2 entrainment (constant rotation number of 0.5). The 1:2 Arnold tongue is bordered by a larger zone of 1:1 entrainment, indicated by the red regions. When stimulation is provided at 130Hz (indicated by the black line), the excitatory population can be entrained at 65Hz for a range of stimulation amplitudes. In all three fitted models, the 1:2 tongue is left leaning (more of the tongue is to the left of the frequency it stems from, see Figs 6A, D and G), which suggests that stimulation frequencies lower than the stem of the tongue are more likely to lead to 1:2 entrainment. The range of stimulation frequencies that can give rise to 1:2 entrainment (given an appropriate stimulation amplitude) is the narrowest in RCS02 (110-160Hz), and the broadest in RCS18 (60-160Hz). In all cases, we note that increasing stimulation amplitude above a certain value will result in the loss of 1:2 entrainment.

**Figure 6:**
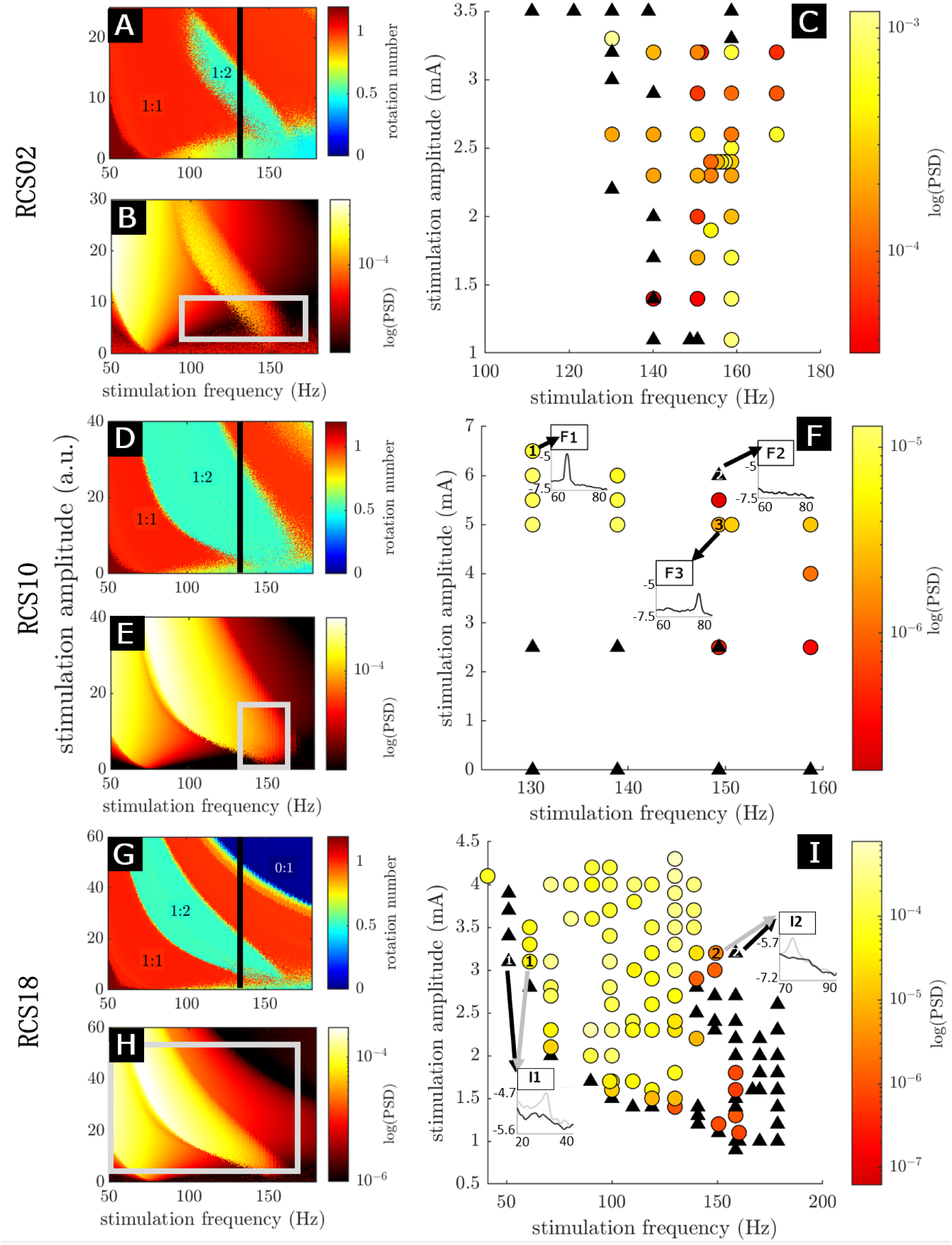
Testing model predictions of a cortical circuit’s response to an external stimulus using human neural data during neurostimulation. Stimulation frequency is the horizontal axis for all panels, while stimulation amplitude is the vertical axis for all panels. Stimulation amplitude has arbitrary units (a.u.) for all model panels (A, B, D, E, G, and H) and is in mA for the data panels (C, F and I). Panels A-C are for RCS02, D-F for RCS10 and G-I for RCS18. Panels A, D and G have colour scales indicating the rotation number (as explained in the *Rotation Number* and *Arnold Tongu*e section) resulting from the stimulation parameters at that point, where 1:1 entrainment is in red and 1:2 entrainment is in green. For these panels, the black line indicates the 130Hz stimulation condition used in [5, 14]. (A, D, G) The entrainment field of the Wilson-Cowan model with the top ranked parameters. The stimulation applied is a single time step pulse with no recharge. (B, E, H) The maximum height of the entrained peaks as predicted by the top-ranked Wilson-Cowan model fit, calculated as laid out in the *Providing Stimulation and Entrainment Analysis in the Model* section. The grey rectangles indicate comparable stimulation parameter ranges to those used in the variable stimulation response data, with stimulation amplitude scaling calculated as outlined in the *Correspondence Between Stimulation Amplitude in the Model and in the Data* section. (C, F, I) The height of entrained peaks obtained from patient recordings for a series of different stimulation parameters. Circles display peak height (represented by the colour scale) of parameters that displayed entrainment (as seen e.g. in inserts F1-2), while black triangles are for parameters that did not display entrainment (as seen e.g. in insert F2). The occurrence of both a black triangle and a circle at the point (140Hz, 1.4mA) in panel C and point (150Hz, 2.5mA) in panel F indicate intermittent entrainment, hence, this will likely be on the boundary of the tongue. Inserts F1-3 and I1-2 show the logarithm of PSDs over frequencies around half the stimulation frequency. For inserts I1-2 the grey arrow and line indicate stimulation parameters where 1:2 entrainment was observed and black where it was not.

The left lean of the 1:2 Arnold tongues does not depend on whether stimulation is applied to the inhibitory population or the excitatory population (see Supplementary Materials section D.5). Additionally, the 1:2 entrainment region exhibits a left lean regardless of whether the stimulation being applied is in the form of a single time step pulse train, pulse trains with various recharge durations or more complicated waveforms (see Supplementary Materials sections D.2 and D.3).

Additionally, the fitted Wilson-Cowan models predict that the highest spectral peaks will occur at the lowest frequencies for which 1:2 entrainment arises, as seen in Figs 6B, E and H.

### Validation of Model Predictions in Human Patient During Chronic Therapeutic Stimulation

The presence of 1:2 entrainment at variable stimulation parameters was investigated in follow-up recordings for the same patients as the Wilson-Cowan model was fitted to (see recording details in the *Data Collection* section). These data were only examined following the core predictions from the model.

### Left lean of the 1:2 tongue, and loss of entrainment for large stimulation amplitudes

For all three patients, the data show regions of stimulation parameters for which 1:2 entrainment can be observed (Figs 6C, F and I), and these Arnold tongues appear to exhibit overall similar shapes to the Wilson-Cowan model predictions (shown in Figs 6A, D and G). For RCS10, while 1:2 entrainment was seen for amplitudes greater than 5.5mA for 140 and 130Hz, it was lost for 150Hz (Fig 6F). This is a robust observation as three repeats lead to the same outcome for this particular combination of stimulation parameters (150Hz, 5.5mA, labelled 2 in Fig 6F). Hence, the 1:2 tongue has a left lean in this set of data with 1:2 entrainment being maintained at higher amplitudes for lower frequencies of stimulation, which is also the case in the model (Fig 6D). RCS18 also exhibits a left-leaning 1:2 tongue. There is no 1:2 entrainment at 2mA for stimulation frequencies above 150Hz, however, in the range of 50-140Hz we can see 1:2 entrainment up to 4mA (Fig 6I). This forms part of the top boundary of the 1:2 Arnold tongue. Additionally, there is a boundary to the right of the 1:2 tongue where no entrainment is seen for any amplitude with stimulation frequencies greater than 155Hz. We are also able to see loss of entrainment on the lower left boundary from 50-150Hz to further demonstrate the left lean. The corresponding model also predicts 1:2 entrainment down to 60Hz in Fig 6G, which is not seen in all other model predictions and is further supported by the data (Fig 6I). Due to patient fatigue, we were unable to chart the right boundary of the 1:2 tongue in RCS02 (Fig 6C). Hence, we are unable to determine if this patient does exhibit a left lean from their data. However, the 1:2 tongue of the model prediction for RCS02 (Fig 6A) is smaller than the other model predictions, with a significantly lower top boundary (stimulation amplitude at which 1:2 entrainment is lost, around 3.5mA). This can also be seen in the corresponding data (Fig 6C).

### Change in entrained gamma power with stimulation frequency and amplitude

Overall, the data validate the model’s prediction of highest power for the lower frequencies within the 1:2 tongue. In RCS10, changing stimulation parameters from 130Hz, 6.5mA to 150Hz, 5mA results in a drop in entrained peak power, as indicated by the colourscale in Fig 6F. The increase in PSD with decreasing stimulation frequency (p *<* 1*×*10*^−^*^4^, Fig 7C) and increasing stimulation amplitude (p *<* 0.05, Fig 7D) are statistically significant. RCS18 also displays a statistically significant increase in entrained peak power as stimulation frequency is decreased (p *<* 0.001, Fig 7E) and stimulation amplitude is increased (p *<* 1*×*10*^−^*^9^, Fig 7F). Power changes with stimulation parameters do not show statistically significant trends within the 1:2 tongue in RCS02 (Figs 7A-B). The increase in entrained peak power with decreasing stimulation frequency in RCS10 and RCS18 is in agreement with model predictions. The increase in entrained peak power with increasing stimulation amplitude is also consistent in the data and the model across the 1:2 tongue, but the relationship reverses in the model when controlling for stimulation frequency (see Fig 6B, E, and H). This is likely to also be the case in the data, although there aren’t enough data points to run a statistical analysis for RCS10.

**Figure 7:**
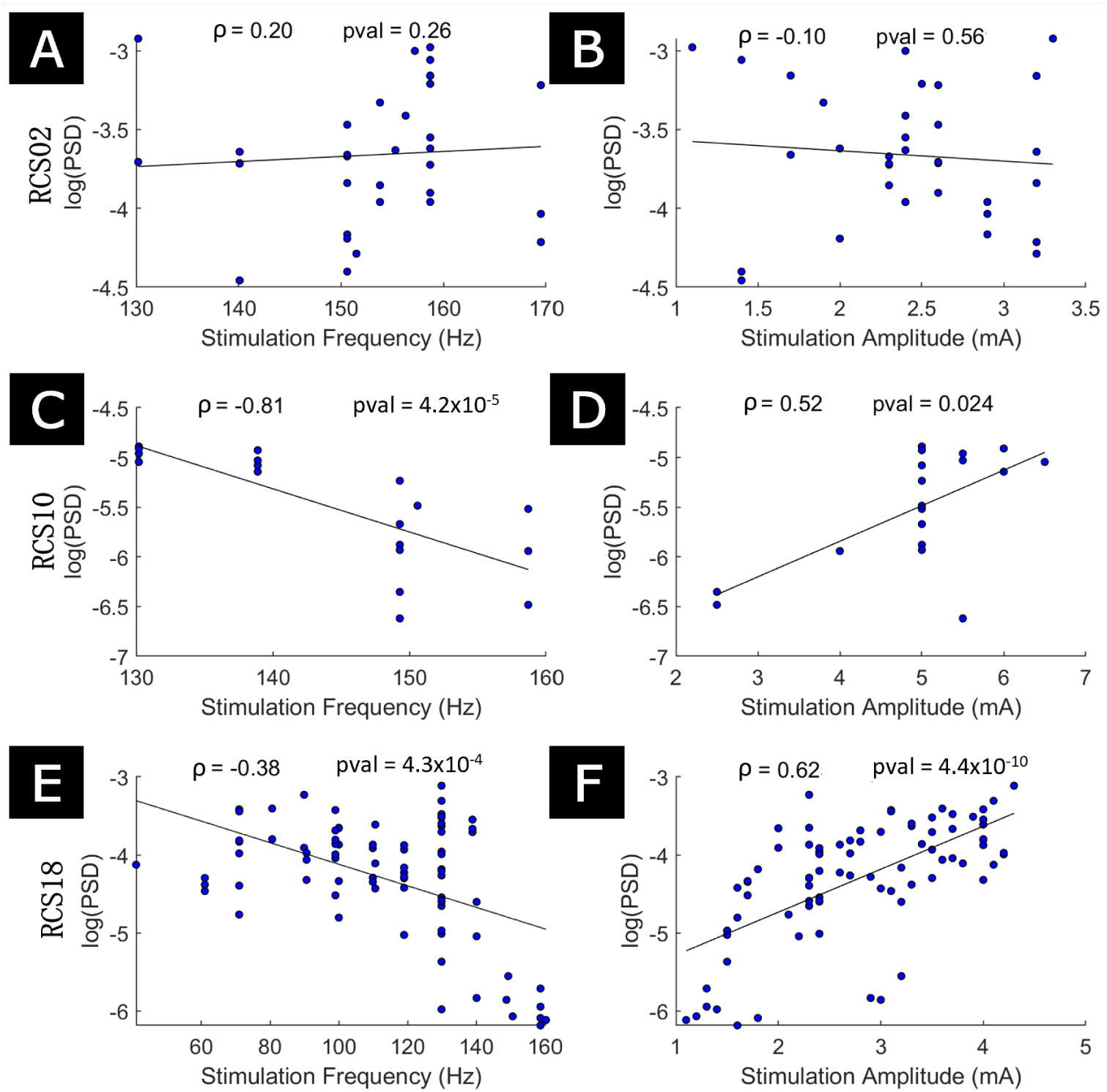
1:2 entrained power correlates with stimulation parameters in two out of three patient datasets. The results for RCS02, RCS10, and RCS18 are presented in the first row, second row, and third row, respectively. The first column display the comparison of the PSD of 1:2 entrained peak heights against stimulation frequency, with stimulation frequency acting as the sole predictor of the linear regression. The second column shows the comparison of the 1:2 entrained PSD against stimulation amplitude, where stimulation amplitude acts as the sole predictor of the linear regression. In all panels, datapoints considered are stimulation parameter combinations that resulted in 1:2 entrainment. The correlations are Spearman’s.

Comparing model predictions of entrained peak power to data provides further support for the models fitted to RCS10 and RCS18 (Fig 8). The correspondence between stimulation amplitude in the model (arbitrary units) and in the data (mA) was established as described in the *Correspondence between stimulation amplitude in the model and in the data* section in the Methods. In Fig 8, green points indicate stimulation parameters resulting in 1:2 entrainment in both model and data (where the tongues overlap). This choice is the most natural to study the relationship between entrained peak power in the model and the data. Black points indicate stimulation parameters that are inside the data 1:2 tongue, but outside the model 1:2 tongue, and thus have an entrained PSD power in the model of very close to zero. This analysis mixes power prediction with tongue shape prediction, but is more conservative which is why we are presenting both results. There is a significant correlation between data and model predictions for RCS10 (p *<* 1*×*10*^−^*^3^, Fig 8B) regardless of whether the model predictions produce 1:2 entrainment or not. RCS18 had a significant correlation for model results that resulted in 1:2 entrainment (p *<* 0.05, Fig 8C), but not for all model predictions. This is due to some stimulation parameters causing 1:2 entrainment far from the natural FTG frequency falling outside of the model 1:2 tongue, where there is a lower PSD (collection of black points in Fig 8C where the model PSD prediction is close to zero). While there were no significant correlations between model and data entrained peak power in RCS02 (Figs 8A-B), the lack of correlation may be explained by a stronger impact on entrainment of fluctuations in the endogeneous FTG in this patient (more details in the discussion).

**Figure 8:**
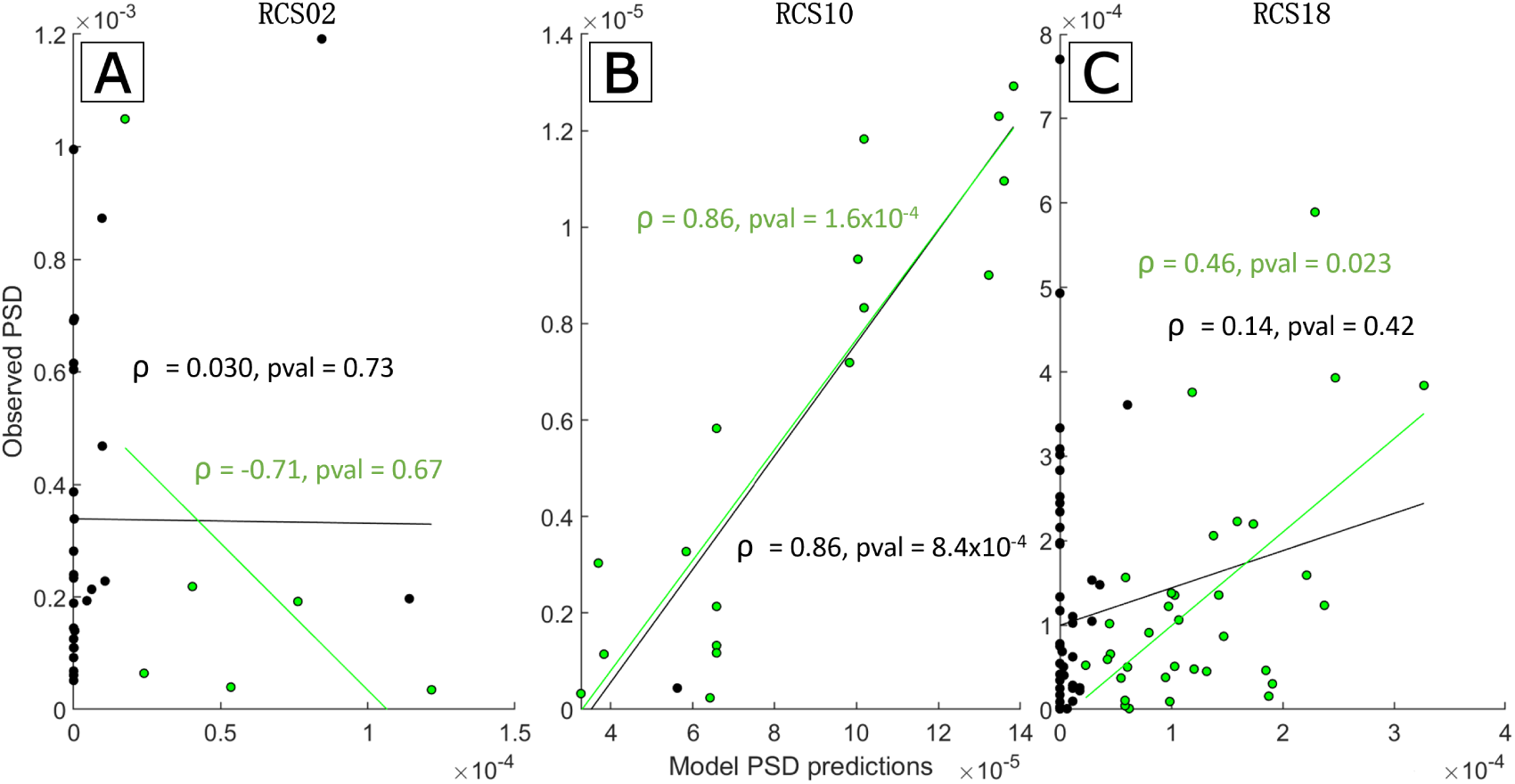
Correlation between model predictions for 1:2 entrained peak power and data. Panel A is for RCS02, B for RCS10 and C for RCS18. Across the three panel, all the data points presented are stimulation parameters that resulted in 1:2 entrainment in the data. The model PSD predictions are obtained using the same stimulation parameters as in the data. Points in green represent stimulation parameters that also achieved 1:2 entrainment in the model, whereas black points did not. The green linear regression fit, *ρ* value and p-value (pval) are calculated using only the points resulting in 1:2 entrainment in the fitted Wilson-Cowan model. The black equivalent are calculated using all points, regardless of whether the fitted model predicted 1:2 entrainment or not. The correlations reported are Spearman’s and p-values are calculated using a permutation test as outlined in the *Correspondence Between Stimulation Amplitude in the Model and in the Data* section.

## Discussion

Here, we studied entrainment of cortical FTG by basal ganglia deep brain stimulation, using a computational model in conjunction with sensorimotor cortex ECoG sensed from chronically implanted bidirectional interfaces in three patients with PD. We show that through fitting a model of interacting neuronal populations to off-stimulation data, we are able to predict the region of stimulation parameters (frequency and amplitude) for which 1:2 entrainment is possible. In particular, our model predicts that 1:2 entrainment is lost when stimulation amplitude is increased above a certain value. Furthermore, the 1:2 Arnold tongue is generally left leaning, implying that 1:2 entrainment can be achieved for stimulation frequencies markedly lower than twice the frequency of the natural gamma rhythm, but not for frequencies markedly higher. Lastly, the model further predicts that there would be a greater entrained gamma power at lower stimulation frequencies that result in 1:2 entrainment. Data recorded during therapeutic neurostimulation, after the modelling results were obtained, showed 1:2 Arnold tongues that generally validate these predictions. Hence, the model can capture a range of sub-harmonic entrainment features without being constrained by entrainment data. The work indicates that effects of DBS at clinically relevant stimulation parameters can be understood using the conceptual framework of a mathematical model representing interacting neural populations being externally perturbed by the periodic neurostimulation.

### Entrainment and adaptive DBS

By solely analysing the presence of 1:2 entrainment, we avoid the prominent artefact at stimulation frequency. Hence, this analysis of the data provides a valuable insight into the neuronal responses to stimulation. Bounding the 1:2 tongue from above, as we lose 1:2 entrainment at increased stimulation amplitudes (all three patients at 150Hz), which provides further evidence that the gamma peak at half stimulation frequency is unlikely to be artefactual. This is aligned with the model prediction that 1:2 entrainment will be lost when amplitude is increased beyond a certain point. Additionally, the model predicts that parameter changes that result in the loss of 1:2 entrainment would see a transition to 1:1 entrainment. However, the presence of 1:1 entrainment is difficult to assess as the resulting power spectral peak can be masked by the stimulation artefact.

1:2 entrainment presents opportunities for adaptive deep brain stimulation (aDBS). aDBS relies on a closed-loop control policy where a biomarker is used to adjust DBS parameters. Cortical gamma-band activity was shown to be a promising candidate biomarker for aDBS [5, 10]. Additionally, entrainment in the gamma frequency band has been linked with dyskinesia [5]. Moreover, 1:2 entrainment has an entirely predictable frequency, which simplifies implementation, particularly for chronic devices with limited spectral recording frequency bands (such as the Medtronic Percept).

### Entrainment in the Wilson-Cowan model

1:2 entrainment is not an intrinsic property of the Wilson-Cowan model (large regions of parameter space do not lead to 1:2 entrainment). Additionally, if the parameters of the Wilson-Cowan model do produce 1:2 entrainment, the 1:2 tongue can also be right leaning or symmetrical about the central frequency. Hence, the parameters of the Wilson-Cowan model need to be tuned to reproduce the data. Among the top-ranked Wilson-Cowan fits, there is some variability between the parameter sets and the corresponding entrainment predictions (see Supplementary Materials section D.1). This demonstrates that the model parameters are non-identifiable. However, as the best fits converge on results that all include a left leaning 1:2 tongue and given the validation of some of the model predictions by follow-up recordings, we can conclude that the fitted model remains a good candidate to make predictions for future investigations. It would be possible to fit Wilson-Cowan model parameters to on-stimulation entrainment data, which may or may not reproduce off-stimulation data. This is not something we are investigating as more value is provided by predicting the response from off-stimulation fits.

While only 1:2 entrainment is investigated here, entrainment will occur at other sub-harmonics of stimulation if there is a neuronal rhythm present to entrain and the corresponding tongue is large enough to encompass the neuronal rhythm. Similarly to 1:2 entrainment, sub-harmonic entrainment at every harmonic of stimulation is not an intrinsic property of Wilson-Cowan models. However, other sub-harmonic entrainment ratios can be observed for certain model parameter sets. By increasing stimulation frequency, for example to around 225Hz, it would be possible to investigate other sub-harmonic entrainment ratios such as 1:3 entrainment of this rhythm.

The observation of the highest spectral peaks occurring at the lowest frequencies of stimulation in the model may be somewhat counter-intuitive, since one could expect more stimulation energy to provide more oscillatory power. However, due to the increased time between successive pulses of stimulation at lower frequencies, the trajectory of the population activity covers a larger distance in phase space (see Supplementary Materials section D.4, specifically Supplementary Materials Fig S.8 for more details on population activity vector fields and trajectories). This means that the range of values that activity reaches for each population is greater, producing a higher power spectral peak for the given resultant frequency. Population activity having a larger range can also be interpreted as there being greater synchrony of neurons within the populations, as increased peak firing rates and decreased minima suggest more neurons are firing together.

Because the endogenous FTG is stronger for RCS02 and stimulation amplitude is lower, more variability in the on-stimulation entrained power is expected, which may explain the worse model performance for this patient (Fig 6 and 8). Specifically, the off-stimulation features of RCS02 (Supplementary Materials Fig S.2A) show a higher gamma peak in the PSD, a larger peak amplitude of the envelope PDF, and a shallower slope at lower amplitudes of the envelope PDF than RCS10 (Fig 4 and RCS18 (Supplementary Materials Fig S.2B). Therefore, it is expected to be more difficult to perturb the underlying oscillators away from the natural gamma frequency of the network, which leads to a 1:2 tongue with a much smaller width over the stimulation frequency axis. Given that the stimulation amplitude used for RCS02 is also smaller than for the other patients (see Fig 6), the strength of the endogenous FTG relative to stimulation amplitude is significantly greater, and natural fluctuations in the endogenous FTG have a larger impact on the power measured. This leads to more variability in Fig 6C and a worse correspondence with the fitted model. However, the fitted model still manages to predict a 1:2 tongue with a smaller area and a significantly lower top boundary compared to the other models, as observed in the entrainment data. Hence, the model is able to predict features of the 1:2 tongue even with stronger endogenous rhythms.

### Study Limitations

As a pilot study, there was no systematic mapping of tongue boundaries, with large regions of untested parameters for some patients and a non-standardised approach to selecting stimulation parameters. Both of these shortcomings will be the focus of further investigations. However, extensive mapping of the 1:2 Arnold tongue boundary may be limited by patient fatigue and discomfort as some parameters tested are subtherapeutic and thus likely to lead to brief exacerbation of motor signs. Additionally, the clinical effects of gamma entrainment were not explored here and will be a focus for future work.

Given that ECoG data represents the activity of populations of neurons, the Wilson-Cowan model (a neural population model) is the appropriate level of description for this type of data. However, this doesn’t allow us to observe or model the behaviour of individual neurons in response to stimulation and during 1:2 entrainment. Our approach is nonetheless adequate to predict stimulation parameters leading to 1:2 entrainment. Additionally, we have not included a population to represent the basal ganglia in our model. This was because there was no consistent subcortical off-stimulation gamma peak to fit a Wilson-Cowan network to for these patients. Subcortical narrowband oscillations in the basal ganglia have been recorded in long term recordings in other patients [10]. It may also be possible to observe 1:2 entrainment in the subcortex and while this study only considers cortical populations, the methodology presented here can be translated to any network of interacting excitatory and inhibitory populations of neurons. Furthermore, we did not consider plasticity in our model. While short-term response to stimulation (such as during the follow-up on-stimulation recordings reported here) can be described by models without plasticity, the several months of chronic stimulation between the prestimulation and follow-up recordings are likely to have led to plasticity, which we did not account for here.

The orthodromic projections from STN or pallidum to cortex are all polysynaptic, and whether the effect of basal ganglia DBS on cortex is excitatory or inhibitory is not well understood. There exist strong connections from STN to GPe [39, 40] and direct GABAergic projections from ChAT (choline acetyltransferase) and prototypic neurons from the GPe to the cortex [40]. Hence, stimulation from these two sites may act primarily on the inhibitory population of the cortex. However, there is not sufficient understanding of the effects of DBS on the basal ganglia as well as the exact nature of the projections from the basal ganglia to the cortex. It is also unclear how antidromic activation of the stimulation targets would effect the motor cortex. To accommodate for this uncertainty, we consider DBS with both GABAergic and glutamergic tendencies (see Supplementary Materials section D.5). Stimulation to our fitted models consistently produces 1:2 entrainment with a left leaning tongue regardless of the population being stimulated. Hence, this choice is inconsequential and the presence of 1:2 entrainment is more specific to the dynamics of the cortical populations than the excitatory bias of the stimulating pulse.

### Implications

This work demonstrates that brain rhythms can have nonlinear responses to stimulation, such as entrainment at harmonics of stimulation frequency, and non-monotonic rhythmic responses to stimulation amplitude. We argue against the simple view that only brain rhythms close to the stimulation frequency can be entrained (through 1:1 entrainment).

These findings might have implications across frequencies. 1:1 entrainment has been reported in the alpha band through single pulse transcranial magnetic stimulation when treating depression [16], with rhythmic visual stimulation [17], and with tACS [41]. If rhythms can lock to harmonics of stimulation frequency, as supported by this study, it is possible that current stimulation protocols targeting any frequency band could induce unexpected responses at sub- or supra-harmonics of the stimulation frequency. Thus, when designing stimulation protocols one should be aware of potential ramifications of stimulation on neuronal rhythms at multiple frequencies. For instance, stimulation targeting lower frequency oscillations, such as beta rhythms, may be able to entrain gamma at a 2:1 rotation number, or even alpha at a 1:2 rotation number. Similar considerations have been employed when designing stimulation protocols in a canine with epilepsy [42]. More recently, a principled approach to selectively promote rhythms close to the stimulation frequency while preventing entrainment at sub- and super-harmonics was put forward in [43].

## Conclusion

We show that for certain network parameters, simple neural circuits can support 1:2 entrainment to DBS. Our fitted Wilson-Cowan model provides theoretical evidence for a neural circuit origin of 1:2 entrainment of cortical FTG oscillation to high-frequency DBS in PD patients. Furthermore, it predicts a larger region of stimulation parameters, at frequencies corresponding to less than twice the natural frequency of the system, for which 1:2 entrainment would be observed. These results are validated by 1:2 Arnold tongue charting from the same patients to whom the model was fitted.

Understanding the variety of effects of stimulation on various brain rhythms would provide valuable insights into designing stimulation protocols to provide maximum therapeutic benefit with minimal side effects. This model provides a first step to predicting these responses. Computational models enable us to experiment with a variety of waveforms without the burdensome tests and validation that would be associated with in patient trials. Prediction of the neuronal responses to stimulation is a fundamental step in the design of future therapeutic protocols. Our model predicts that these responses are not a simple one-for-one mapping of stimulation frequency and amplitudes to brain network activity and that stimulation may have significant effects, even when the stimulation frequency is outside of the frequency band of interest.

## Supporting information

### Supplementary Materials

Further details of the optimisation process, as well as the off-stimulation data and features from the other two patients, the fitting robustness, entrainment predictions for different stimulation patterns, and phase-locking of entrained data signals, are presented here.

## Acknowledgements

The authors are thankful to Huiling Tan for providing helpful comments on the manuscript. We would like to acknowledge the use of the University of Oxford Advanced Research Computing (ARC) facility in carrying out this work. http://dx.doi.org/10.5281/zenodo.22558.

## Data Availability

The data will be made available before publication.

## CRediT Author Statement

**J. Anso:** Conceptualisation, Methodology, Data Curation, Writing - review and editing. **R. Bogacz:** Conceptualisation, Funding Acquisition, Supervision, Writing - review and editing. **S. Cernera:** Data Curation, Investigation, Writing - original draft, Writing - review and editing. **T. Denison:** Conceptualisation, Funding Acquisition, Supervision, Writing - review and editing. **B. Duchet:** Conceptualisation, Investigation, Methodology, Supervision, Writing - review and editing. **S. Little:** Writing - review and editing. **M. Olaru:** Conceptualisation, Data Curation, Investigation, Writing - original draft, Writing - review and editing. **J.J. Sermon:** Conceptualisation, Investigation, Methodology, Validation, Visualisation, Writing - original draft, Writing - review and editing. **M. Shcherbakova:** Data Curation, Investigation, Writing - original draft, Writing - review and editing. **P.A. Starr:** Conceptualisation, Funding Acquisition, Resources, Supervision, Writing - review and editing.

## Supplementary Material

## A Optimisation Details

### A.1 Correlation and Redundancy Between Features

To see consistent population dynamics and entrainment predictions it is required to fit the Wilson-Cowan model parameters to all three features mentioned in the *Fitting Process* section.

We demonstrated this by also fitting to each of the three features individually (approximately 1000 local optimisations per feature). The costs of these optimisations, along with the costs from the optimisation of all three features, are plotted in Fig S.1. In each panel, the trade-off between pairs of features is represented by the Pareto front. As expected, optimising only one feature results in low costs for this feature and significant variability for the other two costs. The results of the optimisation to all three features may not have costs as low as the fits to an individual feature, but they are consistently in the bottom most left corner of the plot and the best trade off between optimising each feature. Moreover, the results of the fits to individual features had greater variability in the entrainment field predictions than the fit to all three features.

**Figure S.1:**
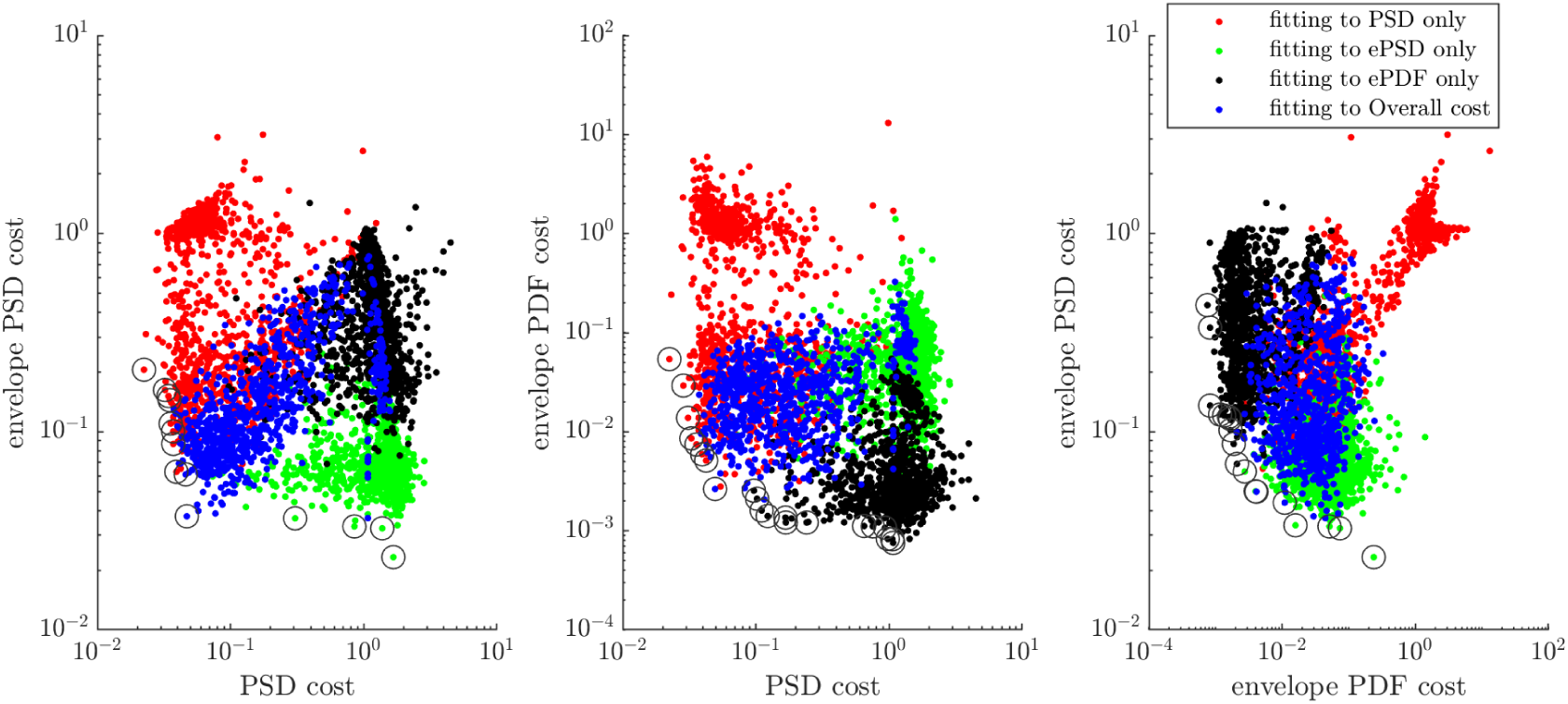
Feature costs when fitting to a single feature only, compared to fitting to all features. Results of fitting to just the PSD (red), envelope PSD (green), envelope PDF (black), and all three features (blue). Panel A shows the comparison of the PSD cost to envelope PSD cost, panel B PSD to envelope PDF cost, and panel C envelope PDF to envelope PSD cost. In all panels, the Pareto front is highlighted by grey rings.

It does not appear that any of these features are redundant to model off-stimulation local field potentials as shown by the low correlation (there is little overlap of costs for the four optimisations in Fig S.1) and the Pareto front in the three panels of Fig S.1. Hence, to capture the full time-domain-based dynamics it is necessary to fit to all three features.

### A.2 Evaluation of Cost

For each parameter set, the model is forward simulated using the Euler-Maruyama scheme for a four second time period, with an additional 0.1 second settling period, and a time step of 10*^−^*^4^ seconds. Following the model simulation, the transient of the settling period is removed and the population activity is z-scored. The three features are then computed in the exact same manner as the data features. The overall cost against the data is calculated by

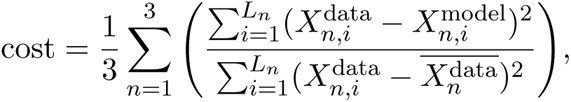

where *n* = [1, 2, 3] for the three features and *L_n_* is the length of each feature, *X_n_*, as in [26]. The optimiser then looks to minimise the cost, which maximises the coefficient of determination, *R*^2^ = 1 *−* cost.

### A.3 Robustness to Noise

Stochasticity introduces the possibility during optimisation that for a given simulation, the randomly generated noise vector may be particularly favourable to minimise the feature costs. However, this parameter set would not represent a local minimum on average. The four second time period of model forward simulation is chosen as a trade off between minimal run time and sufficient individual simulations to minimise the effect of variability due to stochasticity on the parameter set *R*^2^. Following the evaluation of the best model parameter fits, each of the top 20 model parameter sets are simulated three additional times and the features calculated with a different random noise vector. This ensures that the optimal fit was not the result of a single, favourable noise vector. In RCS10 for example, the six top parameter sets featured in S1 D.1 all maintained an *R*^2^ value above 0.92 across the three additional simulations.

### A.4 Top-Ranked Wilson-Cowan Model Parameter Set

Parameters of the top-ranked Wilson-Cowan model fit are given in Table A. The top-ranked parameter set from RCS18 had a smaller, but wider off stimulation gamma peak making the final parameter ranking more prone to variation from noise (see Fig S.2B1). This introduce more variability in the entrainment predictions, but the 1:2 tongue was still consistently predicted to be left leaning in the top 20 parameter sets.

**Table A:** Parameter set from the best ranked fits. The given parameter sets had an average *R*^2^ value of 0.961, PSD cost of 0.038, envelope PSD cost of 0.072 and PDF cost of 0.008 for RCS02, *R*^2^ value of 0.944, PSD cost of 0.052, envelope PSD cost of 0.070 and PDF cost of 0.046 for RCS10, *R*^2^ value of 0.868, PSD cost of 0.156, envelope PSD cost of 0.221 and PDF cost of 0.019 for RCS18, across the 50 simulations lasting 100 seconds each.

| Parameter | $\omega_{EE}$ | $\omega_{EI}$ | $\omega_{IE}$ | $\tau_E$ | $\tau_I$ | $\eta_E$ | $\eta_I$ | $\beta$ | $\zeta$ |
| --- | --- | --- | --- | --- | --- | --- | --- | --- | --- |
| RCS02 | 4.795 | 7.224 | 4.913 | $6.424 \cdot 10^{-3}$ | $4.853 \cdot 10^{-3}$ | 1.977 | -1.020 | 2.216 | 0.087 |
| RCS10 | 1.130 | 1.722 | 3.311 | $9.322 \cdot 10^{-3}$ | $7.329 \cdot 10^{-3}$ | 3.065 | 0.709 | 9.273 | 0.048 |
| RCS18 | 2.590 | 7.679 | 7.140 | $8.595 \cdot 10^{-3}$ | $1.131 \cdot 10^{-2}$ | 4.007 | -0.436 | 3.329 | 0.038 |

## B Off-Stimulation Data and Model Features in RCS18 and RCS02

Off-stimulation data features (PSD, envelope PSD and envelope PDF) are shown for RCS02 and RCS18 in Fig S.2. The features of the top ranked models fits are compared to data features for RCS02 and RCS18 in Fig S.3.

**Figure S.2:**
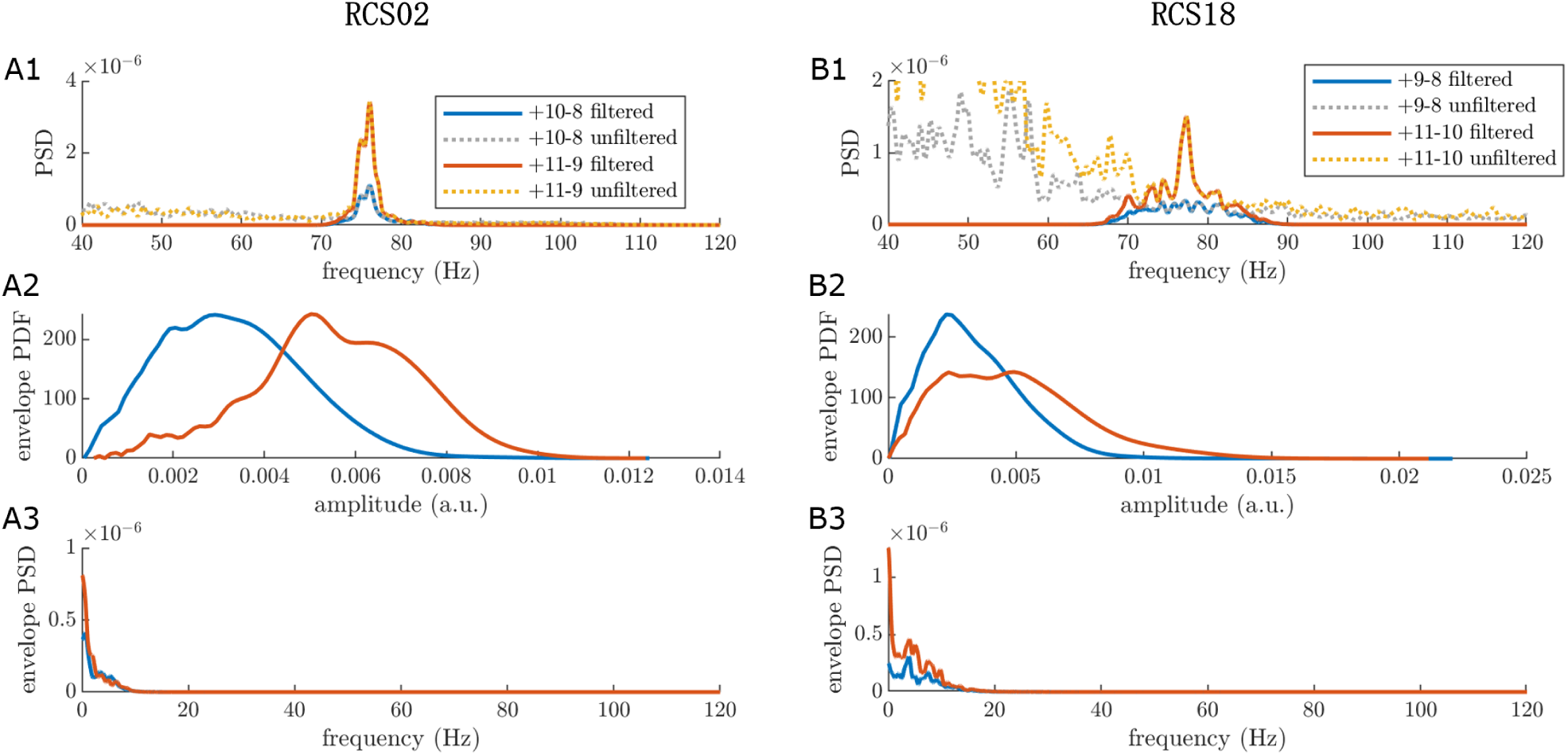
Features of prestimulation human cortical recordings to fit Wilson-Cowan model parameters in RCS02 and RCS18. Features are obtained as described in the main text. The three data features are PSD (1), envelope PDF (2) and envelope PSD (3), for the selected epoch. Panels A1-3 are calculated from RCS02 cortical data filtered between 72 and 78Hz, as the gamma peak occurred at 75Hz. The fitting process for RCS02 was performed with the features from the 11-9 contact as the objective. Panels B1-3 are calculated from RCS18 cortical data filtered between 75 and 81Hz, as the gamma peak occurred at 78Hz. The fitting process for RCS18 was performed with the features from the 11-10 contact as the objective. The solid orange and blue lines display the band-pass filtered cortical signals. The yellow and grey dotted line in the PSD plot.

**Figure S.3:**
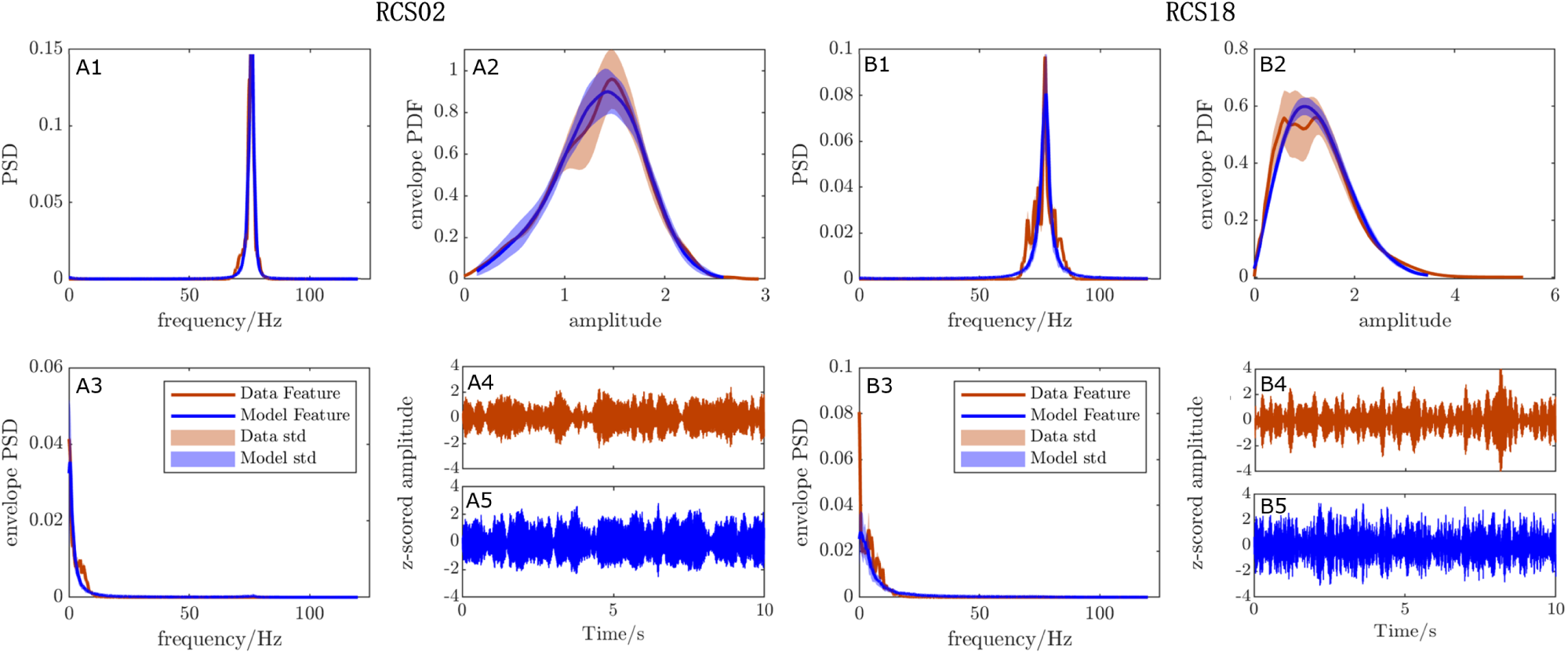
Comparison of the cortical FTG features for z-scored data and the features from the z-scored model output of the best ranked model parameter set for RCS02 and RCS18. Panels A1-5 are the features for RCS02 and Panels B1-5 are the features for RCS18. *R*^2^ = 0.961 for RCS02 and *R*^2^ = 0.868 for RCS18 on average across 50 simulations lasting 100 seconds each. The features are PSD (1), envelope PDF (2), and envelope PSD (3). (4 and 5) Comparison of the band-passed, z-scored, off-stimulation time series from patient data (4) and model output (5) show that the model is capable of replicating similar time-domain dynamics.

## c Sine Circle Map

The sine circle map is the simplest model that describes the influence of periodic stimulation on an oscillator and can provide a first level description of gamma entrainment during 130Hz stimulation. The model stroboscopically observes the phase, *θ*, of a single oscillator of natural frequency *f*_0_, periodically stimulated at frequency *f_s_* and stimulation intensity, A*_s_*. The map between the oscillator phase right after stimulation pulse *i* and its phase right after stimulation pulse *i* + 1 is given by

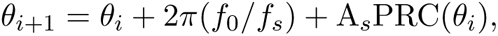

where PRC denotes the oscillator phase response curve and describes the change in the oscillator phase as a function of the stimulation phase. For the sine circle map, the PRC is given by PRC(*θ*) = sin(*θ*). By simulating the sine circle map, we are able to observe a 1:2 Arnold tongue (Fig S.4), which predicts 1:2 entrainment for a 75Hz oscillator at 130Hz stimulation. There exists a specific range of stimulation amplitudes for which we would expect to see 1:2 entrainment of the oscillator at a resultant frequency of 65Hz. This is in agreement with the observations by Swann et al. [5, 14] and provides theoretical grounds for expecting 1:2 entrainment during high-frequency stimulation.

**Figure S.4:**
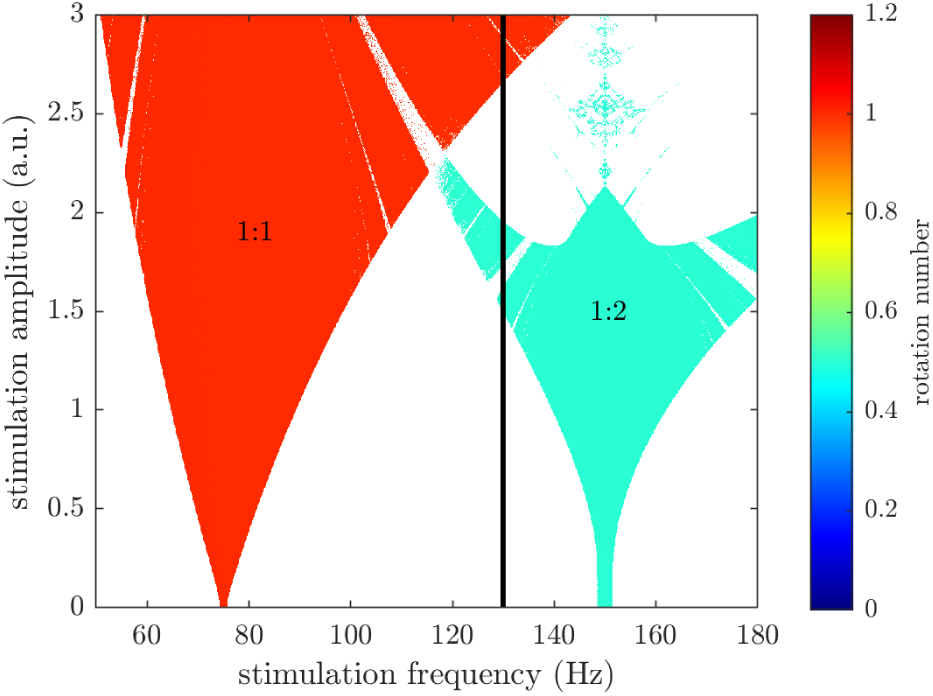
The sine circle map entrainment field. Variable stimulation frequency with a fixed natural frequency of 75Hz. Only showing the 1:1 and 1:2 tongues for clarity, other higher order tongues are hidden.

While the sine circle map can provide a first level description of gamma entrainment, its simplicity results in significant limitations. Firstly, as the oscillator stays on the unit circle, there is no variable amplitude of oscillations. This makes anything more than analysis of a single neuronal unit unreliable. Secondly, the sine circle map only represents a single oscillator. Therefore it is difficult to draw comparisons to ECoG signals that arise from interacting populations of neurons. Thirdly, it is known that pulse shape impacts entrainment behaviour; however, as the sine circle map is stroboscopic, realistic pulses cannot be used. Hence, a model which captures the interaction of neurons, is representative of larger neuronal populations and for which realistic pulse shapes can be used would be more suitable. We therefore consider an interacting neural populations model that is fitted to patient data to predict stimulation parameters that lead to 1:2 entrainment (See the *Wilson-Cowan Model* section in the main text).

## D Fitting Robustness

### D.1 Consistency Across Best Parameter Fits

The left lean of the tongue over the normalised stimulation amplitude axis is a consistent feature across the top model parameter fits, example for RCS10 is shown in Fig S.5.

**Figure S.5:**
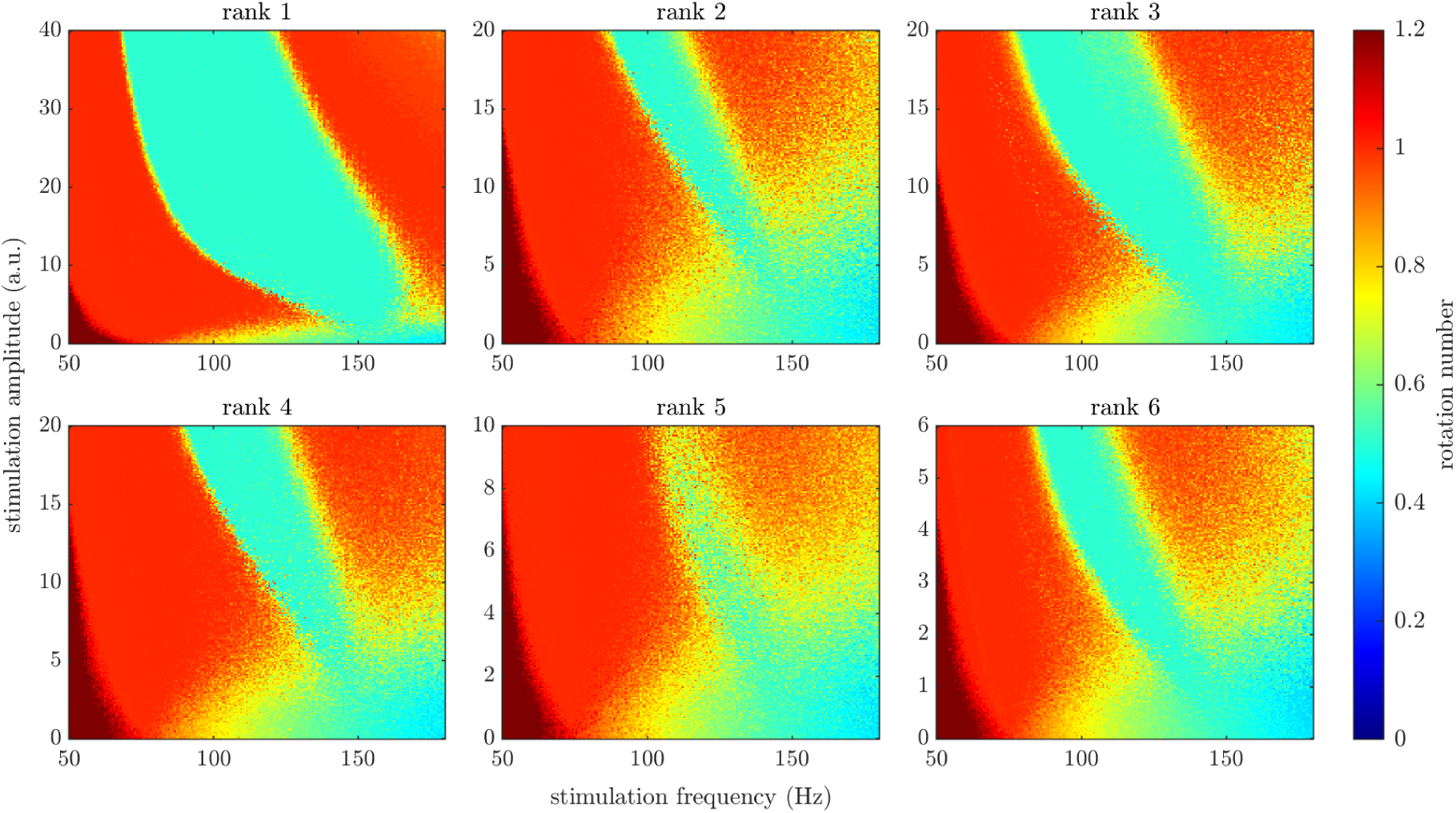
The entrainment fields for the top six ranked parameter fits of the Wilson-Cowan model to RCS10. All model parameter fits presented here had an *R*^2^ value greater than 0.92. Rank five’s entrainment field is observed over a smaller range of stimulation parameters due to the smaller and noisy 1:2 tongue.

### D.2 Alternate Stimulation Pulse Shapes with Recharge All Predict Left Leaning 1:2 Tongues

Throughout this study, we have used the simplest pulse shape with no recharge. However, to achieve charge balancing at the site of stimulation, DBS requires a recharge component following the initial pulse. This component can be active when recharge is a set proportion of the stimulation period, or passive when charge is allowed to flow from the stimulation site via the electrode. The amplitude of the charge balancing component is calculated depending on the frequency of the stimulation and the amplitude of the positive pulse, so that the positive pulse and the negative charge balance have equal areas.

Across pulses with various recharge duration, the 1:2 tongue is present and exhibits a left lean as shown in Fig S.6 (example for RCS10). Despite these similarities, the tongues are present across different amplitude ranges. When recharge is present, the shortest recharge lengths require the greatest amplitude to observe the same extent of the 1:2 tongue. This is attributed to the immediate recharge cancelling the effect of the preceding pulse in the model.

Neurons are likely to respond to hyperpolarising and depolarising stimulation in different capacities. Immediately after a depolarising stimulation pulse, it is likely that a high proportion of the affected neurons will fire. The following hyperpolarising recharge will simply reduce membrane potential during the refractory period. Because of this asymmetrical effect and for simplicity, we chose to use a stimulation pulse with no recharge in this study. However, the prediction of a left leaning 1:2 tongue holds for pulses with recharge of various durations as shown in Fig S.6.

If the model does approximately capture the response of neurons to stimulation pulses with recharge, we would expect that passive recharge or recharge over a longer window would achieve entrainment at lower stimulation amplitudes. Entrainment at lower stimulation amplitudes would potentially increase the efficiency of stimulation protocols that are designed to entrain brain oscillations and minimise the side effects experienced from high stimulation amplitudes.

**Figure S.6:**
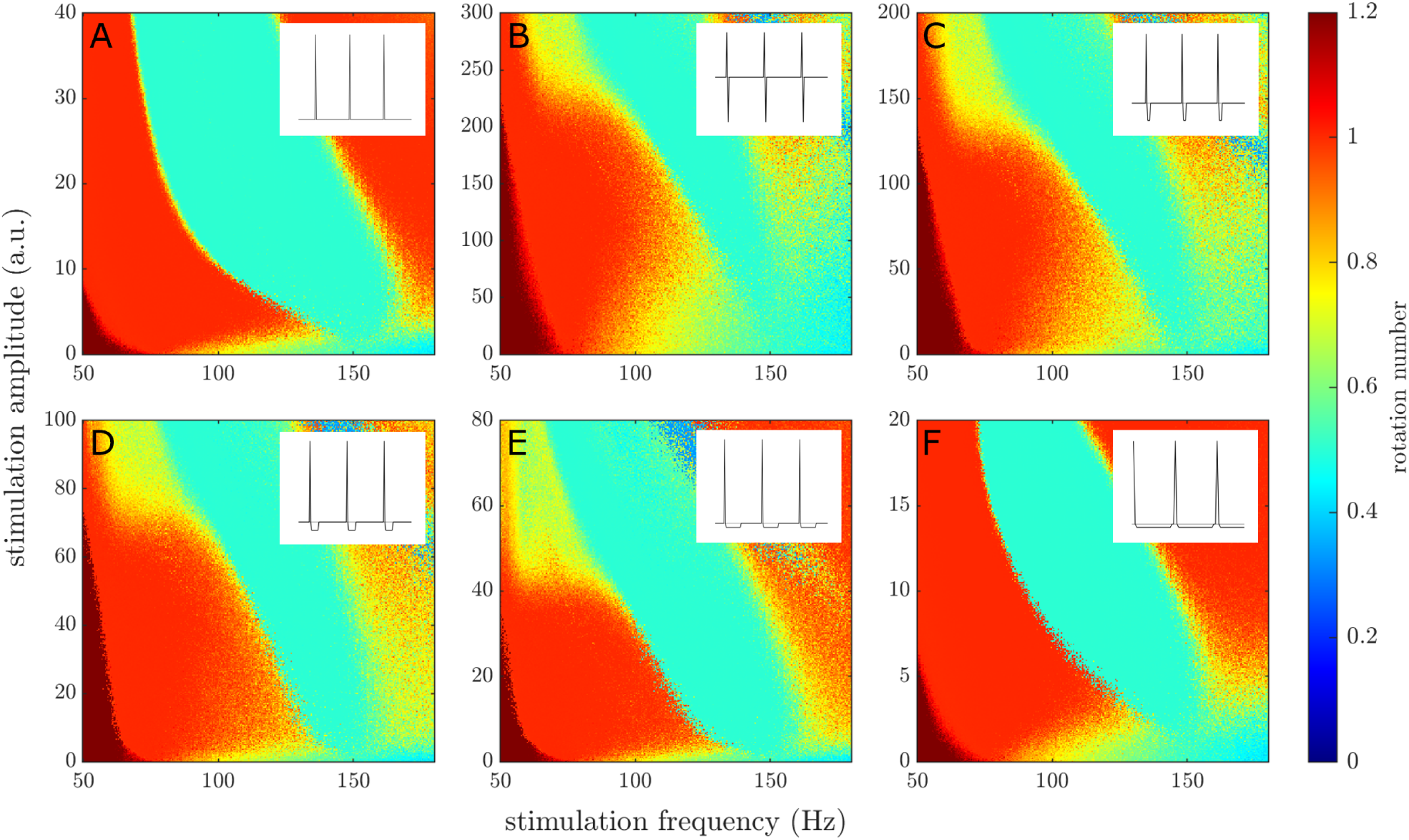
A comparison of six different stimulation pulse recharges applied to the inhibitory population of the top ranked Wilson-Cowan model fitted to RCS10. The panels are categorised as a pulse with no recharge (A), which is the pulse used throughout the majority of the study, 100*µs* recharge (B), 400*µs* recharge (C), 1*ms* recharge (D), 2*ms* recharge (E), and whole period recharge (F), which can be used to approximate passive recharge. The inserts represent a couple of the periods of the corresponding stimulation pulse and the grey line indicates an amplitude of zero.

### D.3 Alternate Stimulation Waveforms Predict Different 1:2 Tongues

As the model is fitted to off-stimulation data, it is possible to apply a variety of stimulation waveforms to either population to predict entrainment responses. Fig S.7 shows the entrainment behaviour for RCS10 at high stimulation frequency for four waveforms: a single time step pulse with no recharge (the same as for Fig 6B), a sine wave, a sawtooth wave, and a square wave. The amplitude scale for each waveform is normalised by the integral of the square of the waveform. This ensures that amplitude levels correspond to the same energy across the waveforms in Fig S.7. The sine wave and the single time step pulse have similar 1:2 tongues at frequencies greater than 150Hz, while the square and sawtooth waves are able to maintain 1:2 entrainment at higher stimulation amplitudes. Additionally, the three non-pulsatile waveforms are able to maintain 1:2 entrainment at lower stimulation frequencies and amplitudes than the pulsatile waveform. This is important to note as a patient may not be able to withstand the higher amplitudes on this scale without experiencing side effects.

### D.4 Tongue Lean and the Wilson-Cowan Vector Field

The left lean of the 1:2 tongue can be explained by analysing deterministic trajectories of the population activities in the E-I activity phase space. Fig S.8 shows that in the 1:2 entrainment region for RCS10, two positive pulses of stimulation occur for each circuit completion of the periodic trajectory (Fig S.8C). As stimulation amplitude and/or frequency are increased, the pulse that occurs at larger values of excitatory activity drops to lower values. Increased frequency (Fig S.8D) means less time for population dynamics to reach more extreme activities. Increased amplitude (Fig S.8A) pushes the activity further away from the fixed point and, hence, requires more time to reach the same value of excitatory population activity.

By considering stimulation parameters along the top right boundary of the 1:2 tongue, these effects can account for the left lean of the tongue. On the boundary, the pulse at higher excitatory population activity of the 1:2 entrained trajectory exactly crosses the fixed point in the absence of noise. In a noisy system it would cross either side of the fixed point, with these cycles corresponding to either 1:2 or 1:1 entrainment. From stimulation parameters on the boundary, changing one stimulation parameter requires mutually changing the other in the opposite direction to maintain this relationship of the higher excitatory activity pulse to the fixed point. For instance, if we increase stimulation frequency and there is less time between the successive positive pulses, we must decrease stimulation amplitude to maintain this relationship. This can be seen in the transition from trajectory A to D in Fig S.8. Hence, this gives us a relationship for the upper bound of the 1:2 tongue where the system transitions to 1:1 entrainment for increasing frequency or amplitude. As this is the upper bound of the tongue, this means that it must be left leaning.

**Figure S.7:**
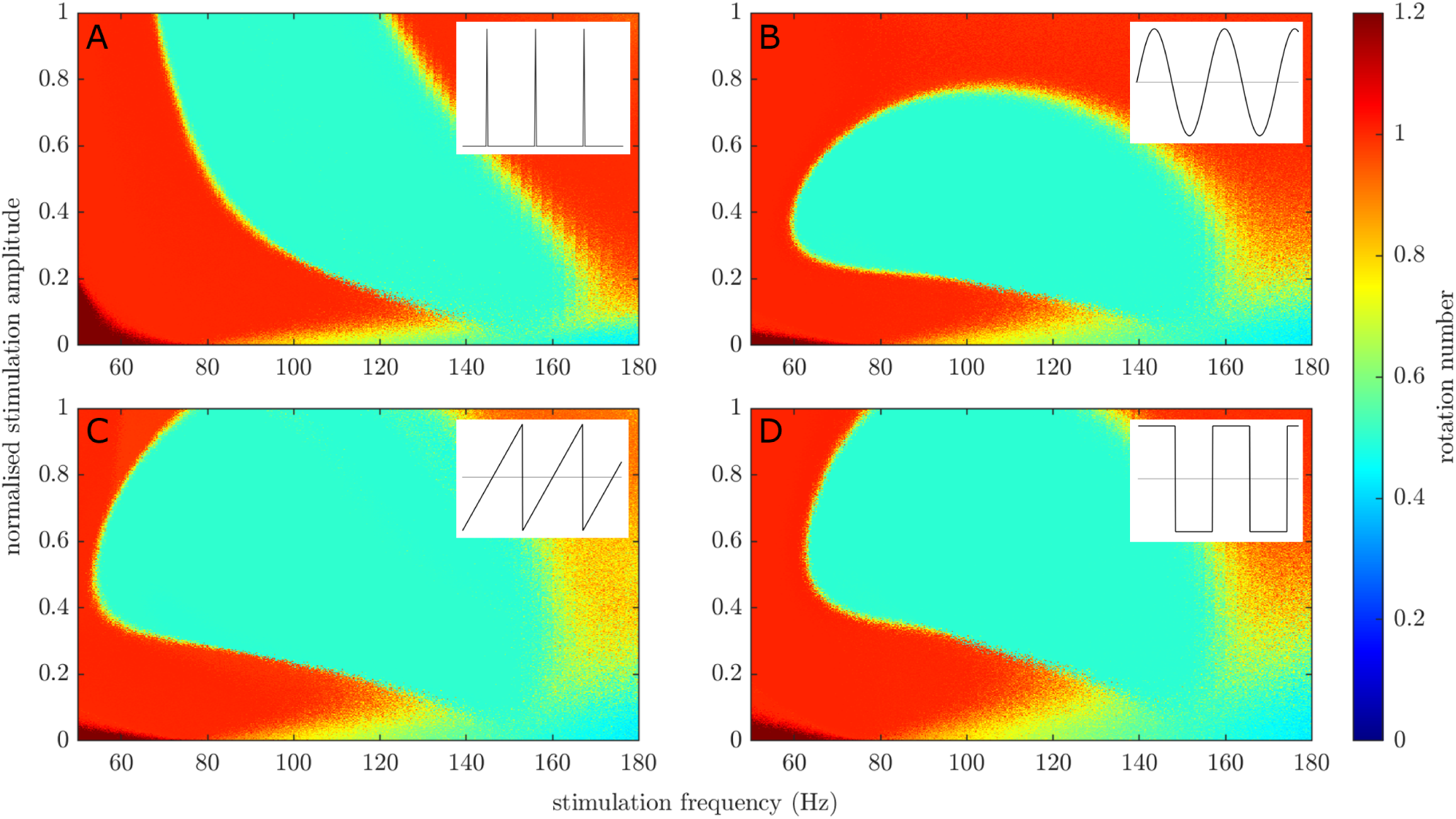
A comparison of four different waveforms of stimulation applied to the top-ranked Wilson-Cowan model fitted to RCS10. In each panel, stimulation takes the form of a single time step pulse with no recharge (A), a sine wave (B), a sawtooth wave (C), and a square wave (D). Stimulation amplitude is normalised against the integral of the square of the waveform. The inserts represent a couple of periods/cycles of the corresponding stimulation waveform and the grey line indicates an amplitude of zero.

The direction of the 1:2 tongue lean depends on the vector field. The entrained trajectory requires more time between pulses to reach the same location within the vector field as stimulation amplitude is increased. The rate of change of population activity is slowed down each period as it consistently reaches darker and slower regions of the colour-scaled angular phase velocity, as shown in the inserts of Fig S.8. If, however, the higher excitatory activity pulse reached a lighter coloured region of increased angular phase velocity, it would require less time between successive pulses to reach the same location within the vector field, i.e. increased stimulation frequency. Therefore, through differences in the vector field, for this hypothetical example, we would predict that increased amplitude would require an increase in frequency to maintain 1:2 entrainment. Unlike the results of our fitting process, this hypothetical example would produce a right leaning 1:2 tongue with a preference for speeding up the natural frequency.

**Figure S.8:**
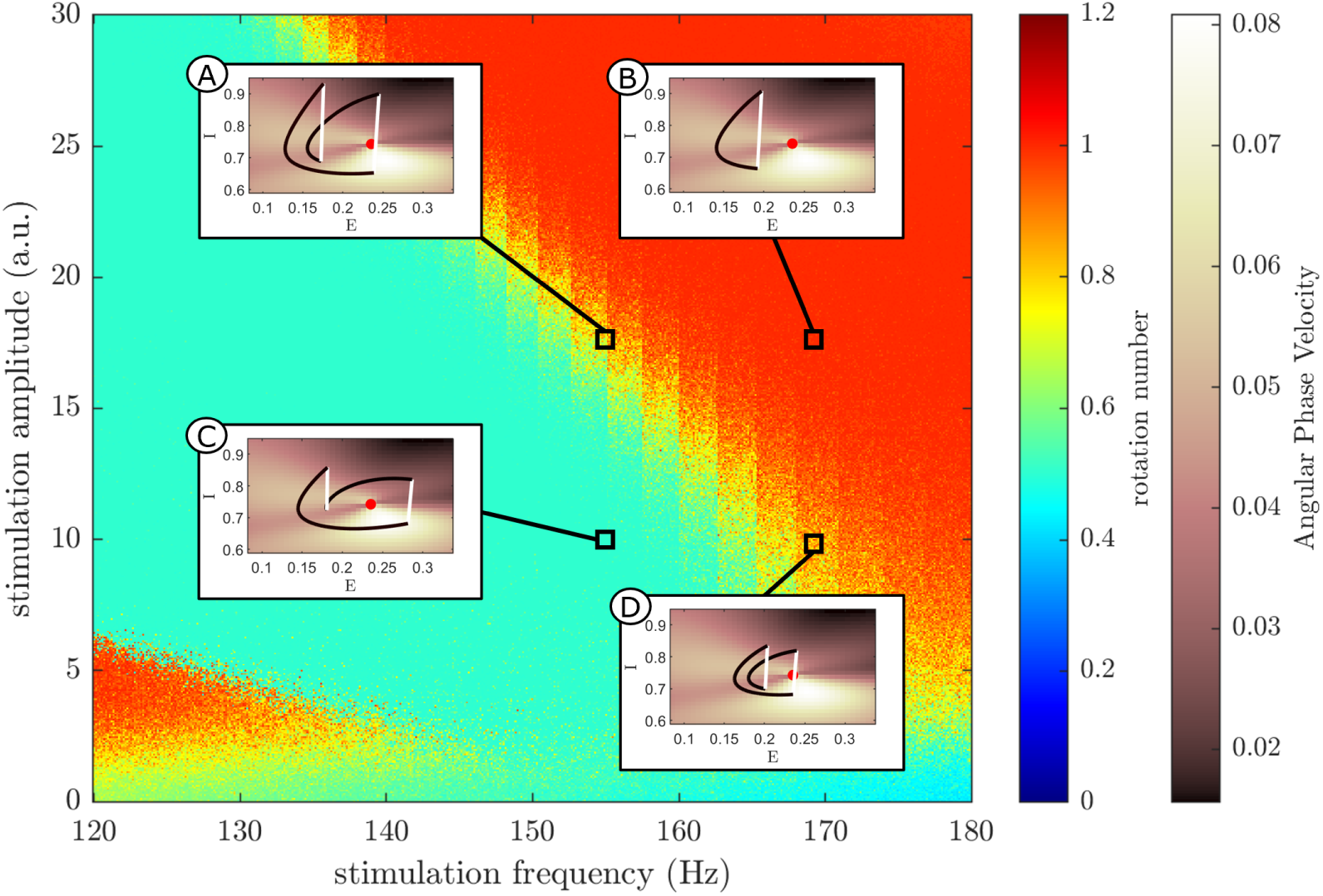
Deterministic trajectories in phase space and the left lean of the 1:2 tongue (example for RCS10). On each of the four trajectory panels, the red dot represents the fixed point for the parameter set. The colourscale of the trajectory is chosen so that white represents a larger jump over an individual time step (following a stimulation pulse) and black is a smaller jump. The angular phase velocity colourbar indicates the magnitude of the vector field, with the colourscale of the entrainment field being identical to Fig 6B. Trajectory A (Amplitude = 18, Frequency = 155Hz) and D (Amplitude = 10, Frequency = 170Hz) show phase space trajectories of stimulation parameters on the 1:2 tongue boundary. Trajectory B (Amplitude = 18, Frequency = 170Hz) lies within the 1:1 tongue and trajectory C (Amplitude = 10, Frequency = 155Hz) is in the 1:2 tongue. The stimulation provided is a single time step pulse with no recharge over the rest of the period.

### D.5 Stimulating the Excitatory Population Instead of the Inhibitory Population

Throughout this paper, we stimulate the models consistently through the inhibitory population as described in the main text. However, the 1:2 tongue is still present when stimulating the excitatory population and while observing the inhibitory population activity (to avoid the population dynamics being dominated by stimulation artefact). Moreover, the key features of the 1:2 tongue are maintained regardless of the population being stimulated. As seen for RCS10 in Fig S.9 and Fig 6D, both 1:2 tongues originate at low amplitude from 150Hz stimulation and have a clear left lean towards lower frequencies. Conversely, stimulation of the excitatory population does not produce 1:2 entrainment at frequencies greater than 150Hz for a single time step pulse with no recharge. 1:2 entrainment above 150Hz was observed in both the model for stimulation of the inhibitory population, in Fig 6D, as well as in the data presented in Fig 6F. Furthermore, the 1:2 tongue resulting from stimulation of the excitatory population transitions to 1:1 entrainment at approximately one quarter the height of the 1:2 tongue of inhibitory stimulation. In the model, 1:2 tongues of similar characteristics originating from both the excitatory and inhibitory population is not a surprise. The network dynamics are responsible for the features of the 1:2 tongue. Therefore a strong periodic stimulus to either population will likely result in the network behaving with a similar response.

**Figure S.9:**
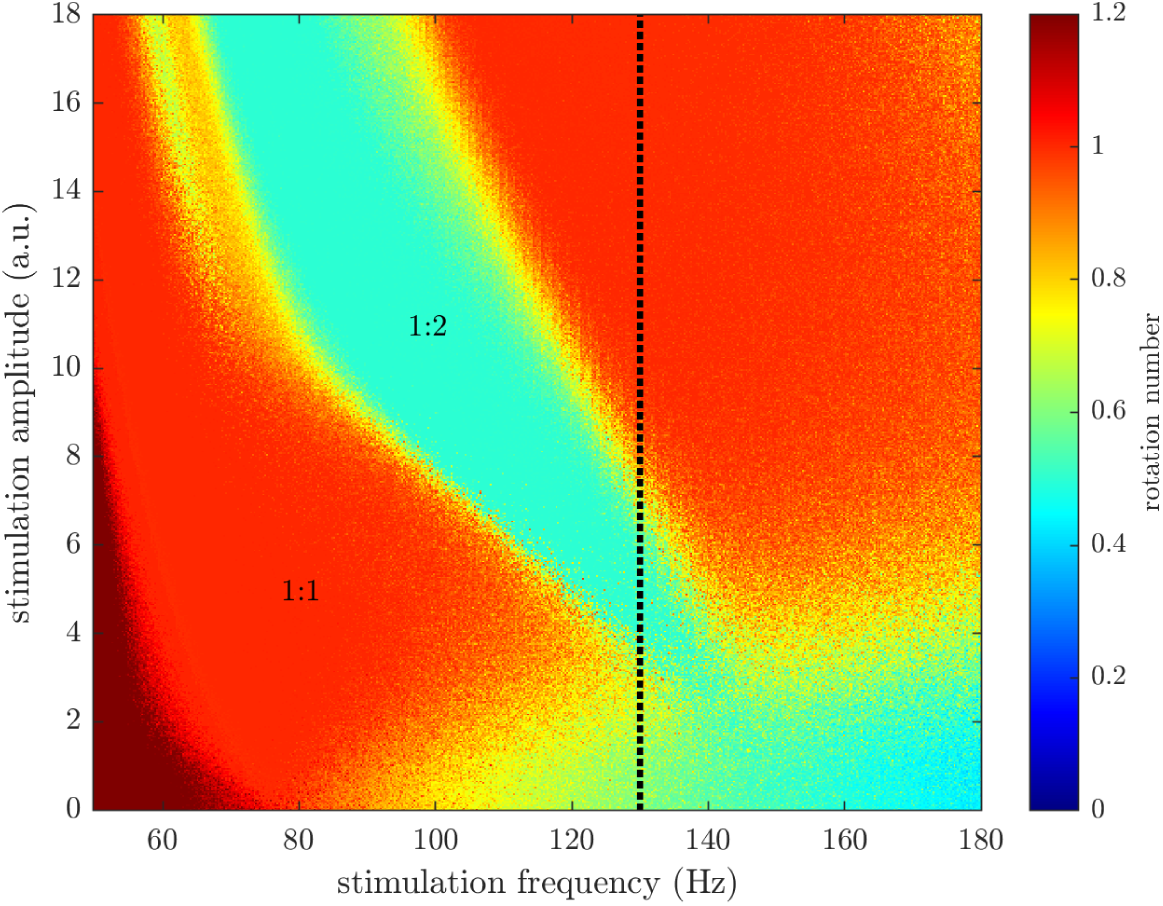
Entrainment field resulting from excitatory stimulation in the Wilson-Cowan model (top-rank fitted to RCS10). Stimulation is for a single time step pulse with no recharge, but applied to the excitatory population and rotation number measured from the inhibitory population. The black dotted line is used to highlight the case of 130Hz stimulation frequency.

### E Entrained Signals Exhibit Phase-Locking

By the definition of entrainment, if oscillators are entrained to a signal then they must exhibit some level of phase-locking. This is observed differently for 1:1 and 1:2 entrainment. For pulsatile stimulation and a 1:1 entrained oscillator, every pulse will occur around the same phase as every cycle of stimulation is matched by a cycle of oscillations. For a 1:2 entrained oscillator, every oscillation cycle experiences two pulses of stimulation. This signal is still phase-locked, but every other pulse occurs around the same phase.

As different modes of entrainment are reflected in the phases at time of stimulation pulses, it is possible to categorise different entrainment regions by plotting these phases. While it is not possible to record from the same contact that is stimulating, it is possible to observe the stimulation artefact from the neighbouring subcortical contacts. By tracking these artefacts and verifying that they align with the stimulating frequency, it is possible to find approximate time stamps of the stimulating pulse train. We record the corresponding phases by taking the Hilbert transform of the cortical signal at these time stamps. Prior to passing the signal through the Hilbert transform, the signals are band-passed between 70Hz and 80Hz for this 149.3Hz stimulation case. This isolates the potentially 1:2 entrained signal, the main observation of interest for this study. These results are plotted in Fig S.10.

**Figure S.10:**
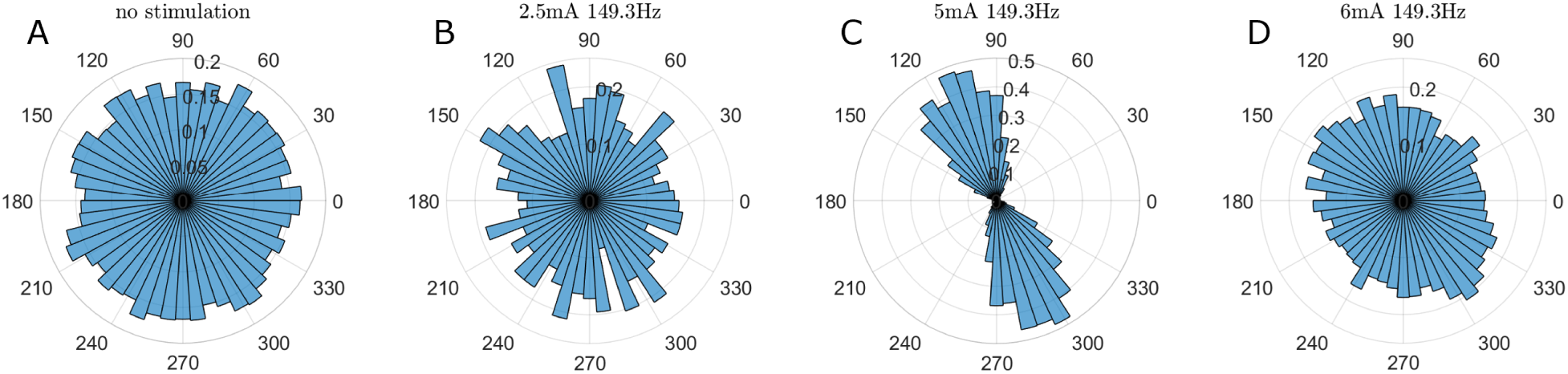
Cortical gamma phases at the time of stimulation for four different stimulation settings in RCS10. Each panel is plotted as a polar histogram and normalised as a probability density function estimate where the sum of the area of the bars equals one. The signals were band-passed between 70 and 80Hz prior to identifying the phases. (A) No stimulation with sham stimulation triggers to record cortical gamma phase at 149.3Hz, (B) 2.5mA for the recording that did not show 1:2 entrainment, (C) 5mA where 1:2 entrainment was observed and (D) 6mA where 1:2 entrainment was not observed.

For Fig S.10A, off stimulation, the recording of phases was triggered by sham stimulation at 149.3Hz. As there is no stimulation, there is no entrainment in panelS.10 A as shown by the absence of preferred phases. Panels S.10B and D did not display 1:2 entrainment in their respective PSDs and do not display preferred phases. In contrast, panel S.10C had a half harmonic peak above the predicted baseline in its PSD and was recorded as seeing 1:2 entrainment. Its corresponding polar histogram displays phases grouped around two, approximately polar opposite, phases. This is in agreement with the spectral evaluation and what we would expect to see from a 1:2 entrained signal.

